# Topography Aware Optimal Transport for Alignment of Spatial Omics Data

**DOI:** 10.1101/2025.04.15.648894

**Authors:** Francesco Ceccarelli, Pietro Liò, Julio Saez-Rodriguez, Sean B. Holden, Jovan Tanevski

## Abstract

Understanding the spatial organization of tissues is essential for uncovering cellular communication, developmental processes, and disease mechanisms. Recent advances in spatial omics technologies have provided unprecedented insight into tissue spatial organization, but challenges remain in aligning spatial slices and integrating complementary single-cell and spatial data. Here, we propose TOAST (Topography-aware Optimal Alignment of Spatially-resolved Tissues), a novel OT-based framework that extends the classical Fused Gromov-Wasserstein (FGW) objective to more comprehensively model the heterogeneity of local molecular interactions. By introducing *spatial coherence*, quantified through the entropy of local neighborhoods, and *neighborhood consistency*, which preserves the expression profiles of neighboring spots, TOAST’s objective function significantly improves the alignment of spatially resolved tissue slices and the mapping between single-cell and spatial data. Through comprehensive evaluations on both simulated and real-world datasets, including human brain cortex Visium data, Axolotl brain Stereo-seq data, mouse embryo seqFISH data, and Imaging Mass Cytometry from multiple cancer types, we demonstrate that our method consistently outperforms traditional FGW and other OT-based alignment methods. Specifically, TOAST improves the accuracy of spatial slice alignment, better preserves cell type compositions, recovers lineage trajectories in developmental brain data, and reconstructs spatial relationships in spatial transcriptomics mouse embryo data. By integrating spatial constraints into OT, our framework provides a principled approach to enhance the biological interpretability of spatially resolved omics data and facilitate multimodal data integration.

## Introduction

Single-cell RNA sequencing enables the profiling of a large portion of the transcriptome at the level of individual cells, and it has profoundly advanced our understanding of disease mechanisms and biological processes ^1^. Recent advancements in spatial omics technologies, particularly in spatial transcriptomics (ST), enable transcriptome-wide gene expression profiling without dissociation, thereby retaining spatial information and offering a complementary perspective on cellular organization. Experimental protocols for generating ST data are well-established and include, for instance, Visium from 10X Genomics ^2^, GeoMx from NanoString ^3^, seqFISH ^4^, MERFISH ^5^, and Slide-seq2^6^. In the context of ST, the ability to align and compare distributions is critical for studying tissues, where different arrangements of cells dynamically interact to maintain tissue function. Changes in cell composition, structure, and spatial organization often underlie the transition from healthy to diseased states ^7^. The alignment of multiple slices is therefore critical for improving the generalization of downstream analyses, establishing plausible trajectories in development, disease progression, or response to perturbation, and integrating multimodal spatially resolved omics data.

Optimal Transport (OT) has become a foundational tool across fields, including economics, machine learning, and biology ^8–12^. By providing a geometry-based approach to realize couplings between two probability distributions, OT facilitates the analysis and alignment of datasets originating from different domains ^13,14^. In the context of tissue biology, OT’s capacity to match distributions in a principled manner makes it an ideal tool for uncovering cellular relationships and dynamics. Recently, OT has been increasingly adopted for tasks involving data integration and alignment. For instance, `PASTE` ^15^ aligns and integrates ST data from multiple adjacent tissue slices, while `PASTE2`^16^ extends `PASTE` to partially overlapped slices. Similarly, `DeST-OT` ^17^ is an OT method for aligning spatiotemporal transcriptomics data, which incorporates cell growth and differentiation objectives in its framework. Related to multislice alignment is the problem of mapping single-cell transcriptomics to spatially resolved imaging/sequencing technologies, hence mitigating the trade-off between lack of spatial information (single-cell technologies) and resolution and gene coverage (spatially resolved technologies). For instance, Rahimi et al. ^18^ proposed `DOT`, a multi-objective optimization framework for transferring features across these data modalities, thus integrating their complementary information. In Klein et al. ^19^, the authors introduced `moscot`, a flexible OT framework that implements widely used OT objectives —Wasserstein, Gromov-Wasserstein, and Fused Gromov-Wasserstein— as well as extensions that relax marginal constraints for unbalanced OT ^8^ and account for cell growth and death. Collectively, these works highlight the versatility of OT in addressing a broad range of challenges in the integration and alignment of single-cell and spatial omics data, particularly through the use of the Fused Gromov-Wasserstein objective.

Despite demonstrating promising results, these methods exhibit key limitations that can compromise both their accuracy and biological interpretability. Specifically, while they leverage OT to infer correspondences between slices by accounting for global spatial organization (intra-sample distances), they do not incorporate the local spatial organization, which plays a crucial role in shaping cellular states. In cancer biology, for instance, the tumor microenvironment is highly structured, with immune cells, fibroblasts, and malignant cells forming spatially distinct niches that regulate immune evasion, therapeutic resistance, and tumor progression ^20,21^. Similarly, in developmental biology, spatial gradients of morphogens and transcriptional programs orchestrate cellular differentiation and tissue patterning, making local spatial context essential for accurately reconstructing lineage trajectories ^22^. Consequently, the omission of domain-specific spatial constraints in OT-based frameworks can lead to biologically implausible mappings that do not capture the true spatial architecture of tissues.

In this work, we introduce TOAST (Topography-aware Optimal Alignment of Spatially-resolved Tissues), a novel topography aware framework for spatial omics, which explicitly incorporates spatial local constraints into the optimal transport objective. TOAST extends the classical FGW formulation by introducing two additional terms, *spatial coherence* and *neighborhood consistency*, to account for spatial organization and molecular heterogeneity. We extensively evaluated TOAST on both simulated and real spatial omics datasets for the tasks of multislice alignment and integration of single-cell and spatial data, demonstrating its advantages over traditional FGW and other alignment methods. Our evaluation is based on data coming from spatial transcriptomics technologies such as 10x Visium, Stereo-seq and seqFISH, and spatial proteomics technologies, such as Imaging Mass Cytometry. By leveraging optimal transport in a spatially informed manner, our method provides a tool for studying tissue organization, tracking cellular dynamics across conditions, and enhancing multimodal data integration.

## Results

### Topography Aware Optimal Transport

A spatial omics dataset is composed of a series of spatially resolved tissue slices, each defined by a tuple 𝒮 = (**X, S**) where **X** ∈ ℕ^*n×p*^ is, for example in the case of transcriptomics, the matrix of *n* spots and *p* genes and **S** ∈ ℕ^*n×*2^ contains the spatial coordinates (*x, y*) of each spot. Given a pair of tissue slices, 𝒮^1^ and 𝒮^2^ from two adjacent tissue sections or time points containing *n*_1_ and *n*_2_ spots respectively, Optimal Transport aims to find a transport matrix 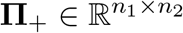, whose positive entry Π_*ij*_ describe the probability of transporting spot *i* in 𝒮^1^ to spot *j* in 𝒮^2^. To this end, several recent works^15,16,19^ proposed the use of the Fused Gromov-Wasserstein distance and minimized the following objective function

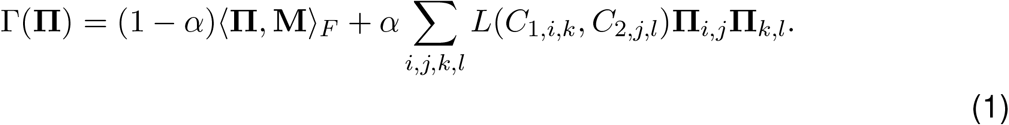

The Frobenius inner product term, ⟨**Π**,**M**⟩ _*F*_, corresponds to the *Wasserstein component*, as it minimizes the pairwise transport cost encoded in the cost matrix **M**. In the context of spatial transcriptomics, the Wasserstein term encourages mappings between spots with similar expression profiles (see Methods). The second term, ∑_*i,j,k,l*_ *L*(*C*_1,*i,k*_, *C*_2,*j,l*_) **Π**_*i,j*_Π_*k,l*_, represents the *Gromov-Wasserstein component*, as it captures the structural dissimilarity between spatial slices. *C*_1,*i,k*_ and *C*_2,*j,l*_ represent the intra-slice spatial distance between spot *i* and *k* and spot *j* and *l* in 𝒮^1^ and 𝒮^2^, respectively. Here, *L*(*C*_1,*i,k*_, *C*_2,*j,l*_) measures the dissimilarity between the local contexts *C*_1_ and *C*_2_, making the OT framework sensitive to relational or structural differences (see Methods). The hyperparameter α ∈ [0, 1] controls the balance between the Wasserstein and Gromov-Wasserstein components.

The state and function of cells within a tissue are affected by interactions with neighboring cells, extracellular matrix components, and local signaling gradients ^23^. The intra and intercellular relationships form distinguishing and representative spatial patterns across tissues and conditions ^24^. Processes such as immune responses, tissue regeneration, and cancer progression often depend on the spatial organization and interaction of different cell types. Furthermore, recent studies on tissue organization in cancer patients have identified the presence of spatially clustered regions, dominated by a single cell state, and disorganized regions, exhibiting a variety of cell states ^25^. As a result, it is pivotal to be able to model these interactions and account for molecular heterogeneity while defining a successful transport plan. To this end, we introduce TOAST (Topography-aware Optimal Alignment of Spatially-resolved Tissues), a generalized topography aware OT-based framework for intra-, intersample, and temporal alignment and mapping of spatially resolved omics data (Figure 1). To decouple the diversity of cell states from the aggregated expression in the immediate neighborhood of each cell, TOAST incorporates two new terms to the objective function in Equation 1.

**Figure 1:**
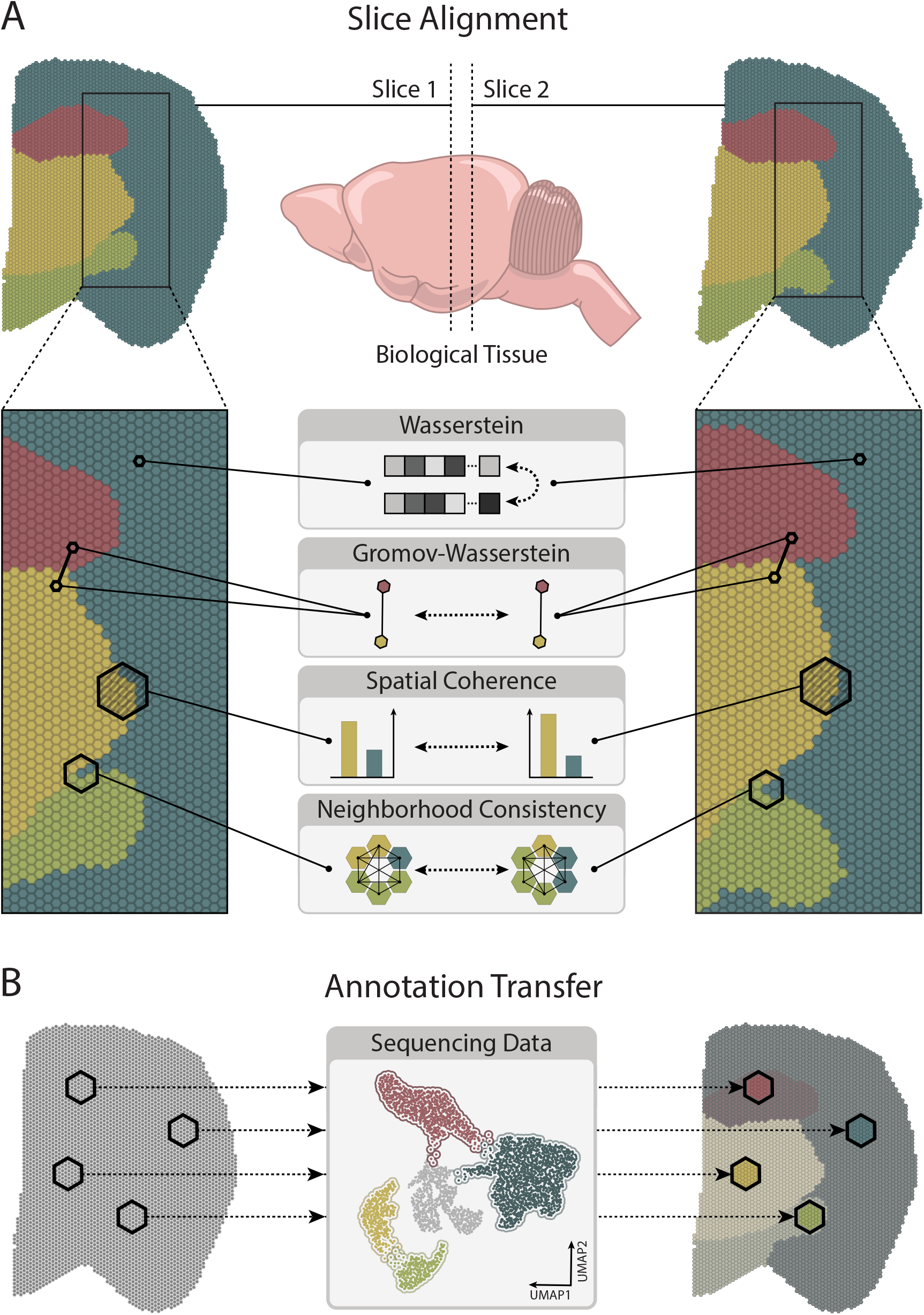
TOAST: a topography aware Fused Gromov-Wasserstein Optimal Transport framework for spatial omics. (A) Slice Alignment: We align spatial tissue slices by incorporating spatial coherence and neighborhood consistency into the optimal transport objective to better preserve local tissue organization and molecular heterogeneity. (B) Annotation Transfer: TOAST enables the integration of multiple slices of spatially resolved omics as well as localization of dissociated cells to spatial locations by mapping cellular states while preserving the spatial context.

First, we introduce a *spatial coherence term*, designed to capture the heterogeneity in the spatial neighborhood of each spot. In detail, using the (*x, y*) coordinates, we construct a spatial graph *G* where each node represents a spot, and edges connect each spot to its closest neighbors. Given a node *i*, we define 𝕡_*i*_(*k*) to be the probability of a neighbor of *i* belonging to class *k ∈K*, where *K* denotes the set of cell labels. We then compute the spatial coherence score for a node *i* as the entropy of the label distribution around *i*:

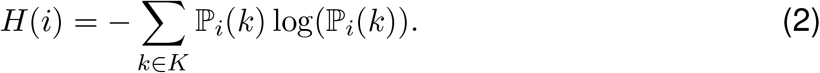

The cell labels typically correspond to manually annotated cell types or cell states, or labels transferred from an available single-cell reference atlas. In the context of the optimization task, however, annotations are not strictly required. Instead, a set of discrete cell labels can be obtained by clustering highly variable genes across tissue slices. *H*(*i*) provides a straightforward interpretation: it quantifies the degree of heterogeneity of a given spot’s neighborhood. A higher *H*(*i*) value indicates a more diverse neighborhood, with multiple cell labels being represented among the neighboring spots. Conversely, a lower *H*(*i*) value suggests a more homogeneous neighborhood, dominated by a single cell label. Incorporating spatial coherence into the OT objective function enables a more refined modeling of spatial interactions. Specifically, it encourages solutions that respect the local heterogeneity of the tissue environment, ensuring that the transport plan aligns with the tissue’s local structure. Following the notation in Equation 1, we define **C**_**3**_ to be the dissimilarity of the spatial entropy scores of spot *i* in 𝒮^1^ and spot *j* in 𝒮^2^ as

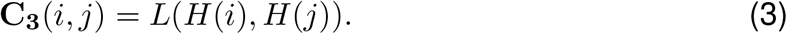

Second, to further enhance the preservation of the spatial context and model the locally consistent intercellular relationship patterns ^24^, we introduce a *neighborhood consistency term*. This additional term is designed to ensure that the average expression profiles of a cell’s neighborhood are preserved across slices. As before, a graph *G* is constructed using the (*x, y*) spatial coordinates. For each node *i* in *G*, we consider its set of neighbors 𝒩 (*i*) and compute the average expression profile across all genes within this neighborhood. We define **C**_**4**_ to be the distance between the average neighborhood expression profiles of spot *i* in 𝒮^1^ and spot *j* in 𝒮^2^ as

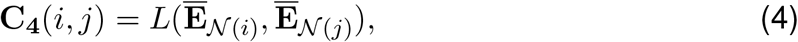

where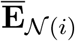 and 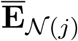 represent the average expression profiles of the neighborhoods of spots *i* and *j*, respectively. The neighborhood consistency acts as a regularization term and penalizes transport plans that map spots exhibiting neighborhoods with drastically different expression profiles, thereby ensuring that the spatial context of each spot is preserved across slices. By integrating both the spatial coherence and neighborhood consistency terms into the classical FGW formulation, we define the objective function of TOAST as

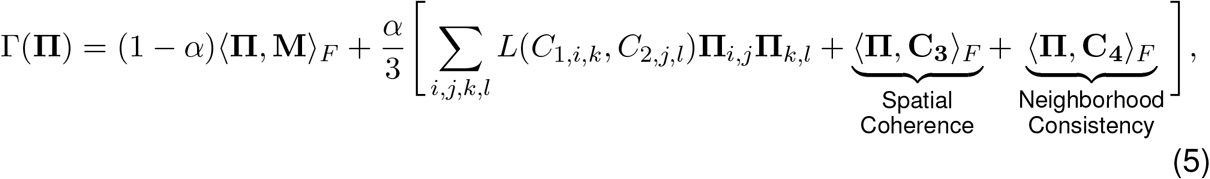

where α ∈ [0, 1] controls the trade-off between the feature similarity term and the equally-weighted structural terms. In practice, an entropy-based regularization term *H*(**Π**) = ∑_*i,j*_**Π** _*i,j*_ log(**Π**_*i,j*_) is added to the optimization problem in Equation 5 to control for smoothness and accelerate the optimization of **Π**^26^. We further constrain the optimization such that the total mass transported from 𝒮^1^ exactly matches the total mass transported to 𝒮^2^ resulting in the balanced OT optimization formulation

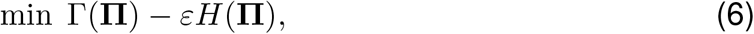

subject to:

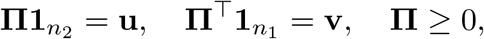

where *H*(**Π**) is the entropy of the transport matrix, *ε >* 0 controls the strength of this entropic regularization, 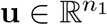 and 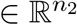 are the mass vectors corresponding to the distributions of spots in 𝒮^1^ and 𝒮^2^, and **1**_*n*_ is a vector of ones.

### Evaluation on simulated data One-dimensional simulation

Following the evaluation scenario presented by Halmos et al. ^17^, we simulated one- and two-dimensional slices with eight-dimensional feature vectors for each spot (see Methods). For the one-dimensional simulation, we defined two tissue slices 𝒮^1^ and 𝒮^2^, each containing 101 spots and two different cell types, A and B. The feature vectors for each cell type were designed to be orthogonal and dependent on the one-dimensional spatial location of each spot ^17^. 𝒮^1^ contained 51 spots of cell type A and 50 spots of cell type B, while 𝒮^2^ had 49 spots of cell type A and 52 of cell type B. In addition, to mimic the organization of real biological tissues, we introduced clustered regions, where only one cell type was present, and disorganized regions, where both cell types were mixed. Figure 2 (A) shows an example of the spatial organization of the simulated one-dimensional ST data. We compared the classical Fused Gromov-Wasserstein (FGW) formulation in Equation 1, with our topography aware formulation, TOAST, that integrates spatial coherence and neighborhood consistency in the objective function as in Equation 5. As evaluation metric, we assessed mapping accuracy as the sum of probabilistic alignment weights **Π**_*ij*_ over all pairs (*i, j*) of spots in the true alignment. For the true alignment, spots were defined as corresponding if they shared the same cell type or were located in regions with the same spatial organization (that is, clustered or disorganized). We defined the former as *cell type accuracy*, and the latter as *coherence accuracy*. For our experiments, we tested multiple *α* values (0.1, 0.3, 0.5, 0.8, 1), progressively increasing the weight of the spatial component. Figure 2 (B) shows the comparison between FGW and TOAST in terms of the above-mentioned metrics. TOAST consistently matches or outperforms FGW in terms of cell type accuracy across all αvalues, indicating its robustness in preserving cell identity. Compared to FGW, our topography aware transport plan preserves the spatial neighborhoods of the slices, as reflected by the coherence accuracy metric. Figure 2 (C) shows a qualitative comparison between the transport maps from FGW and TOAST. In TOAST’s alignment, connections in disorganized regions are notably stronger and better mapped, reflecting the model’s ability to preserve the local context of spatially disorganized areas. In contrast, FGW yields weaker and less consistent connections, as it primarily minimizes transport and structural costs without accounting for the molecular heterogeneity or the neighborhood context of the cells.

**Figure 2:**
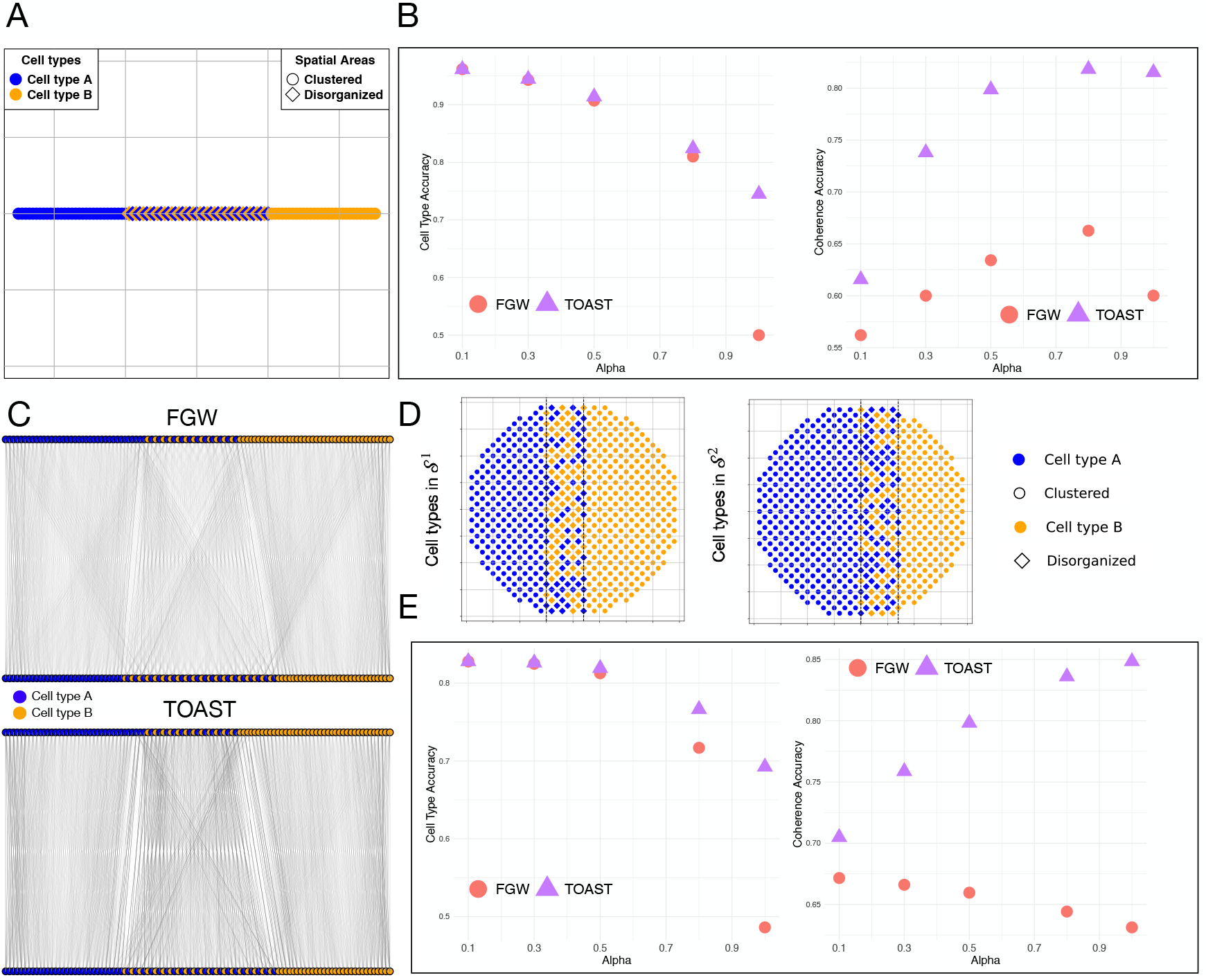
Evaluation on one- and two-dimensional simulated spatial slices. (A) Example of the spatial cell type distribution for slice 𝒮^1^ for the one-dimensional simulation. (B) Accuracy comparison between the traditional FGW formulation and our topography aware formulation, TOAST, in terms of cell type and spatial coherence for the one-dimensional simulated ST data. (C) Qualitative comparison between the transport maps from FGW and TOAST. (D) Spatial cell type distributions for slice 𝒮^1^ and 𝒮^2^ for the two-dimensional case. (E) Accuracy comparison between FGW and TOAST in terms of cell type and spatial coherence for the two-dimensional simulated ST data.

### Two-dimensional simulation

Similar to the one-dimensional case, we simulated ST data in two dimensions (see Methods). Both slices contained 624 spots, with 𝒮^1^ being composed of 278 spots of cell type A and 346 spots of cell type B, and 𝒮^2^ of 385 spots of cell type A and 239 spots of cell type B. As in the one-dimensional case, clustered and disorganized areas were defined (Figure 2 (D)). Figure 2 (E) shows the transport map accuracy for the two-dimensional case, comparing FGW and TOAST across different values of α. On both cell type and coherence accuracy, TOAST consistently outperforms the traditional FGW formulation. For small values of *α*, the cell type mapping performance remains similar as the spatial information is not fully exploited. As *α* increases, FGW performance sharply declines compared to TOAST. At larger *α* values, FGW fails to correctly map cells to spatial regions, whereas TOAST better captures local context and preserves spatial coherence.

### Intra-, intersample and temporal alignment

#### Human dorsolateral prefrontal cortex Visium

To assess the ability of our framework to map spots across slices in a real dataset, we analyzed a human dorsolateral prefrontal cortex (DLPFC) data generated using 10x Visium ^2^. This study sequenced 12 tissue slices spanning six neuronal layers plus white matter (WM) from the DLPFC in three human brains. Manual annotations of the tissue layers were provided in the original study. One of the samples from this dataset is shown in Figure 3 (A). We assessed the pairwise alignment of all consecutive slices for each sample and compared the accuracy, measured by the number of correctly matched cell annotation pairs across the slices, between FWG (Equation 1), TOAST (Equation 5), DeST-OT^17^, Tangram ^27^, Paste2^15,16^ and NovoSparc ^28^. Figure 3 (B) presents the results of this comparison. By explicitly modeling neighborhood composition and spatial expression, TOAST outperforms the other methods across all pairwise alignments and samples. TOAST’s superior performance stems from its ability to explicitly model local cell type composition within the tissue. Given that the brain cortex is a highly layered and spatially organized structure (Figure 3 (A)), where distinct neuronal and glial cell populations are arranged in well-defined regions, a transport plan incorporating neighborhood consistency and spatial coherence is able to yield more biologically meaningful mappings. By preserving the local spatial organization of spots, TOAST ensures that cell type transitions align with the inherent structural properties of the cortex, leading to more accurate cross-slice alignment. While DeST-OT and FGW also show good accuracy, Tangram and NovoSparc fail to align spatial slices accurately in most comparisons, leading to the lowest scores in Figure 3 (B). Unsurprisingly, Paste2 reaches similar results to FGW, as they share the same objective function. We further explored the ability of the models to reconstruct the local spatial heterogeneity by comparing the cell type distributions between cells in the source slice and cells in the aligned slice by quantifying the Jessen-Shannon divergence (JSD) of their nearest 20 neighbors. A lower JSD value signifies a closer resemblance in cell type distribution between predicted and true cell neighbors. Figure 3 (C) shows that TOAST better preserves the spatial cell type composition compared to other methods, consistently reaching lower JSD scores. Along with alignment of consecutive slices, we designed a more challenging set of experiments involving alignment of non-consecutive slices from the same brain, and cross-sample alignment (a slice from one sample is pairwise aligned with a slice from another sample). Figure 3 (D) shows the accuracy results of this comparison. As in the consecutive alignment experiments, Tangram and NovoSparc achieve the lowest accuracy, with DeST-OT and FGW exhibiting higher accuracy scores. However, as illustrated in the boxplots and measured by the Wilcoxon rank-sum test, TOAST outperforms all the other methods with a statistically significant margin across all three experiments. Finally, our topography aware framework also exhibits the strongest performances when comparing the average accuracy and JSD for each cell annotation across all three experiments (Figures S1–S3).

**Figure 3:**
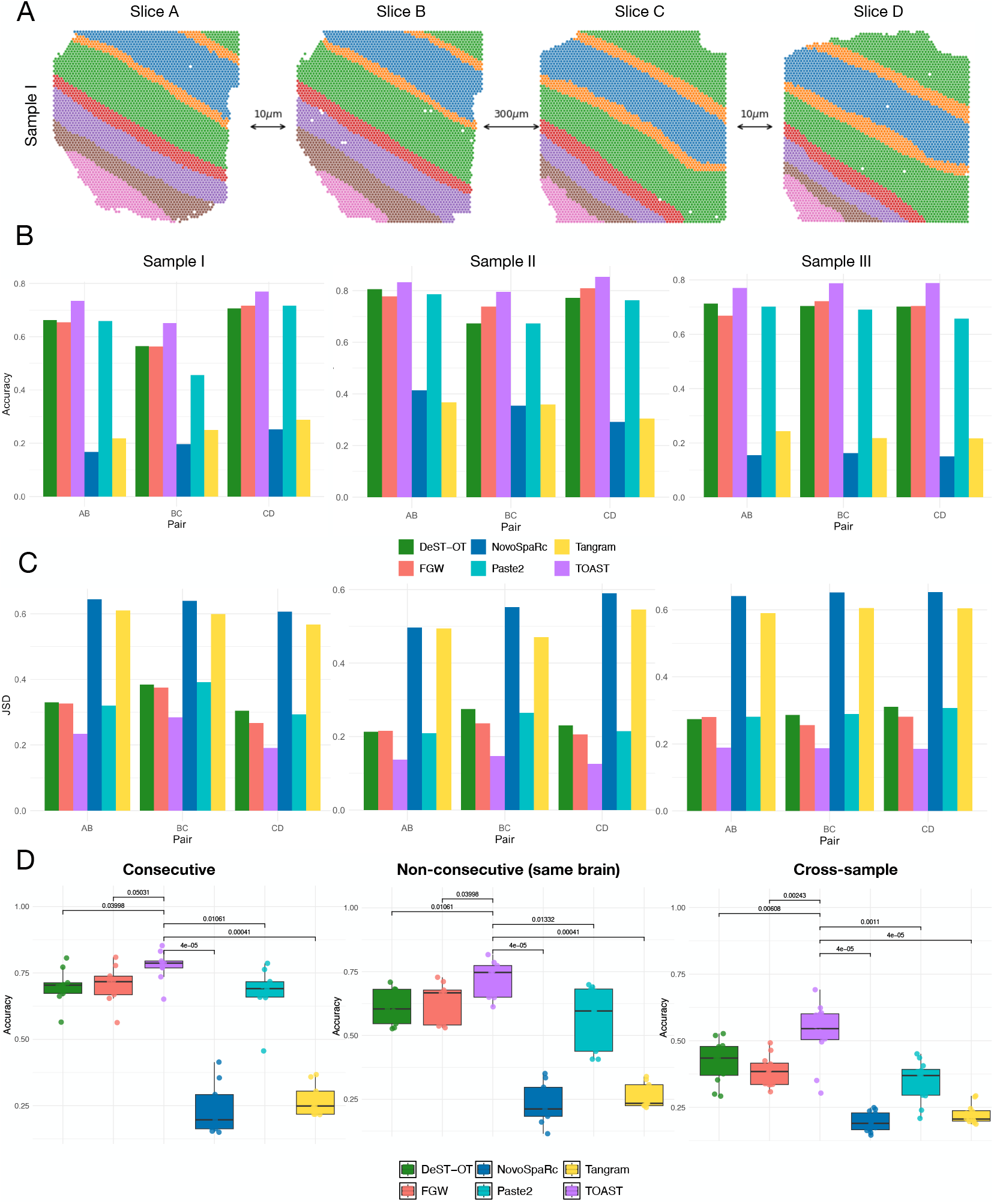
Evaluation on the human dorsolateral prefrontal cortex (DLPFC) data. (A) One sample of DLPFC with four slices where spots are colored according to the original annotations from Maynard et al. ^2^. (B-C) Quantitative comparison of the pairwise alignment of all consecutive slices for FWG, TOAST, DeST-OT ^17^, Tangram ^27^, Paste2^15,16^ and NovoSparc ^28^ in terms of accuracy and Jessen-Shannon divergence (JSD), respectively. (D) Accuracy results for all methods on consecutive slices, non-consecutive slices from the same brain, and cross-sample alignments.

#### Axolotl brain Stereo-seq

In Wei et al. ^22^, Stereo-seq was used to measure gene expression in the telencephalon, a brain region in the Axolotl (*Ambystoma mexicanum*), a species of salamander, at different developmental time points. Figure 4 (A) depicts the spatial organization of cells across the five developmental stages (Stage 44, Stage 54, State 57, Juvenile and Adult) colored by cell type. As before, we assessed the pairwise alignment of all consecutive slices and compared TOAST against the best-performing method from the previous section, DeST-OT. Figure 4 (B) shows that across all values of *α*, by explicitly taking into account spatial neighborhood information, TOAST consistently reaches the highest accuracy. Although the improvement margin is smaller than in the well-layered human dorsolateral prefrontal cortex, TOAST is still capable of outperforming DeST-OT in all alignments. In Figure 4 (C), we quantify the models’ ability to preserve cell type composition across alignments. TOAST exhibits lower JSD scores in all pairwise alignments, indicating that it better preserves the composition of the local spatial neighborhood compared to DeST-OT, which shows greater deviation from the original cell type distributions (performances for each individual cell type are shown in Figure S4). We then examined the cell type transition matrix produced by TOAST for each pair of consecutive time points. From this matrix, we depict the most frequent transitions, supported by at least 50 occurrences, in Figure 4 (D). Consistent with findings in Halmos et al. ^17^, we observed that immature cell types expressing early developmental markers disappear after the juvenile stage, where immature cell types like common myeloid progenitor (CMP) or medium spiny neurons (MSN) transition into their respective mature cell types. Developmental ependymoglial cells (dEGCs) disappear at Stage 57, with TOAST predicting their primary transition into ribEGCs, which reside in the ventricular zone—the same spatial region as dEGCs. Finally, a directed cycle between cckIN and MSN emerges, particularly during the juvenile-to-adult transition, likely due to their co-localization in the striatum region of the brain.

**Figure 4:**
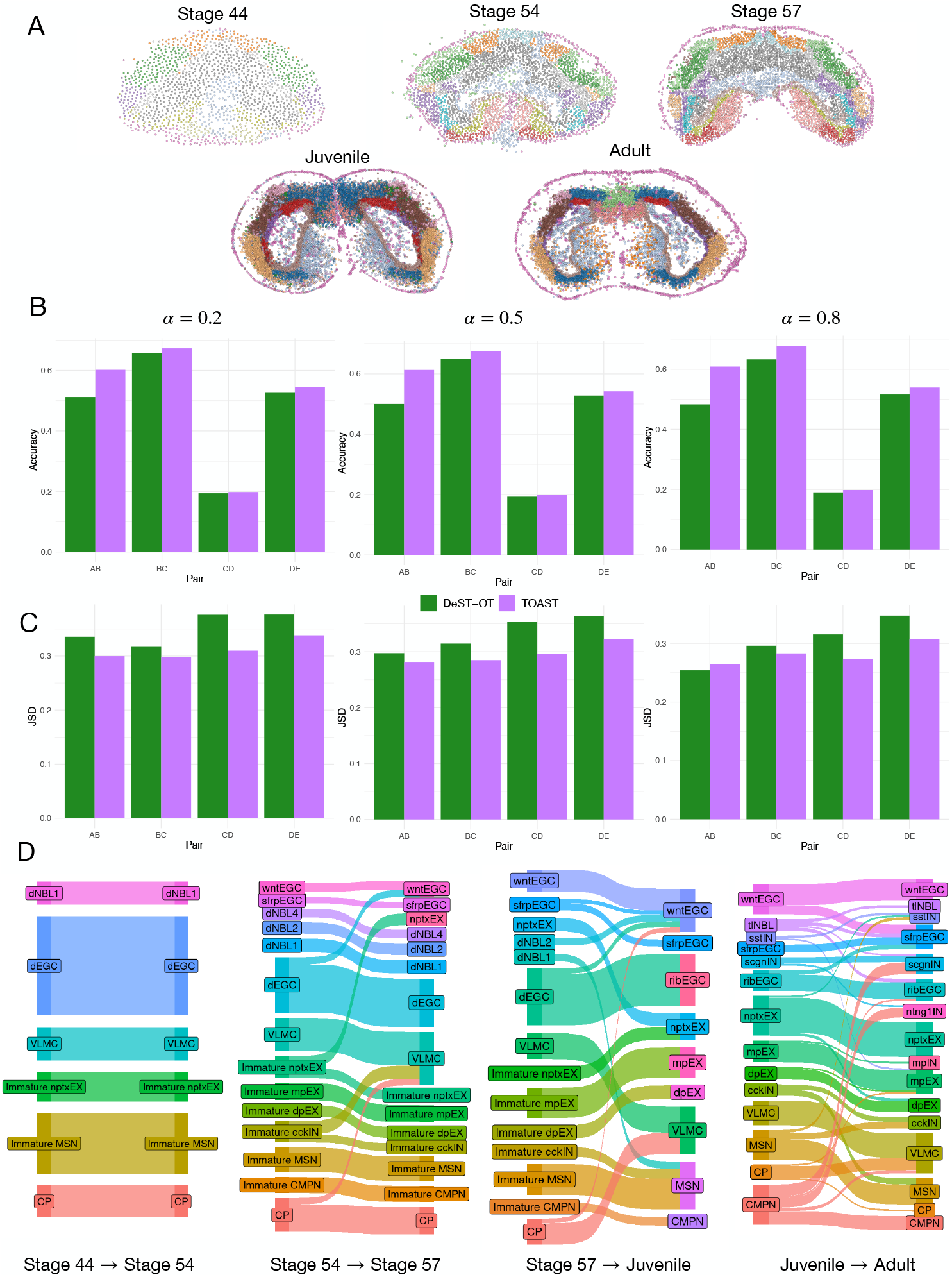
TOAST recovers developmental trajectories in Stereo-seq Axolotl brain. (A) Stereo-seq at different developmental time points colored by cell types. (B-C) Quantitative comparison of the pairwise alignment of consecutive slices for TOAST and DeST-OT ^17^ in terms of accuracy and Jessen-Shannon divergence (JSD), respectively. (D) Most frequent cell type transitions predicted by TOAST for each pair of consecutive developmental stages. Our framework accurately captures the development of immature cells into their respective mature cell types.

#### Imaging Mass Cytometry of different cancer types

While all experiments so far focused on transcriptomics data, our OT framework finds application also on other types of spatial omics. As an example, we considered Imaging Mass Cytometry (IMC) from sections of 4 patients with squamous cell carcinoma of head and neck, breast cancer, non-small cell lung cancer and colorectal cancer respectively from the Integrated iMMUnoprofiling of large adaptive CANcer patient cohorts ^29^ (IMMUcan). Single-cell measurements for 40 proteins, along with annotations of major immune and cancer cell types, were available for each section. Figure 5 (A) shows four sections, annotated by the cell type and by the expression of three markers (CD8, CD68, CDH1). We aligned all intra-sample slices, resulting in a total of 18 comparisons, and used the cell type annotations to measure the accuracy of DeST-OT and our topography aware formulation, TOAST. Figure 5 (B) quantifies the accuracy of both methods across all alignments.

**Figure 5:**
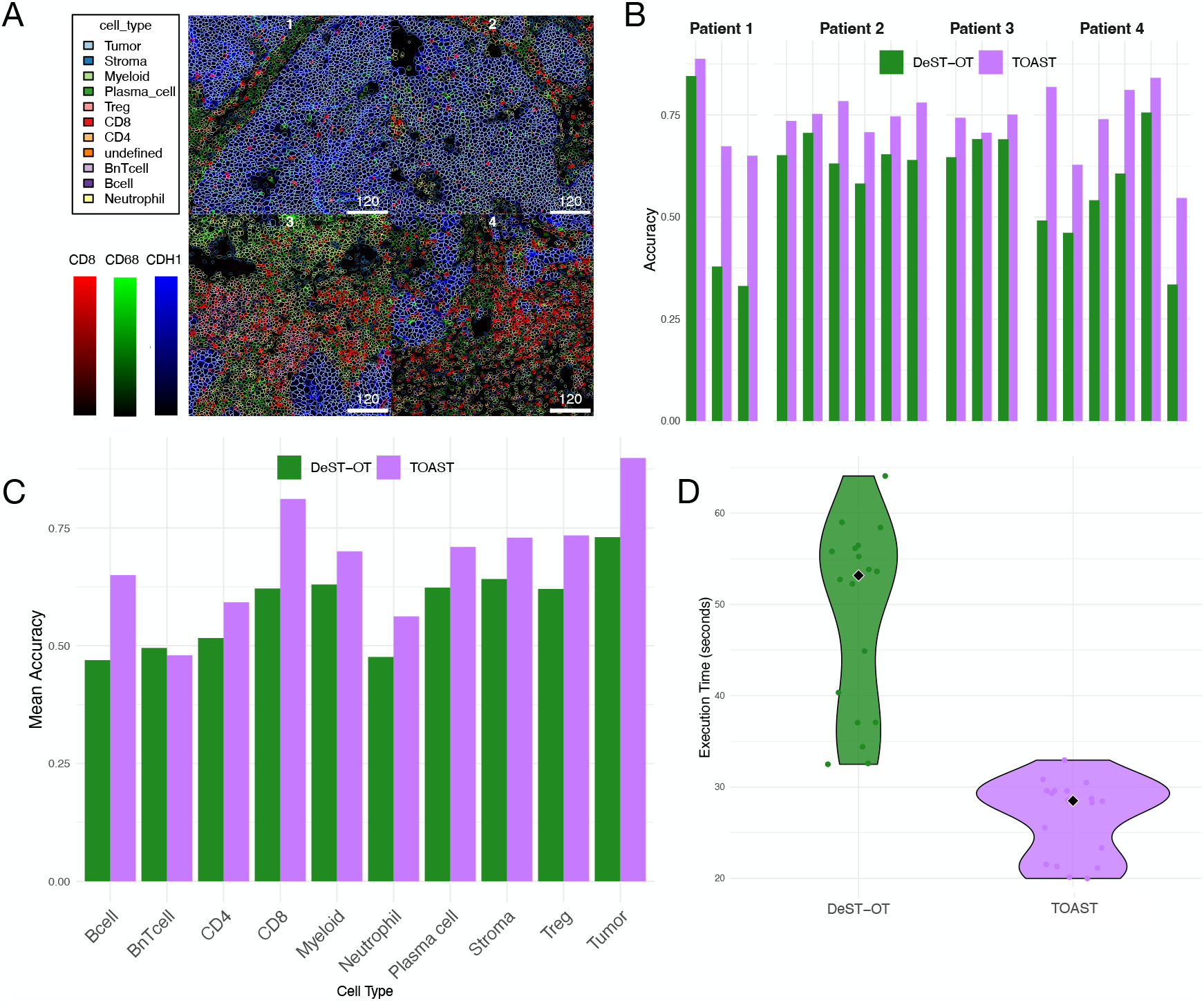
TOAST accurately aligns Imaging Mass Cytometry data of different cancer types. (A) For four example sections from IMMUcan ^29^, the spatial distribution of different cell types is shown, with colors representing CD8a, CD68, and CDH1 protein expression. The outlined cells correspond to different annotated cell types. (B) Accuracy for TOAST and DeST-OT ^17^ for all intra-sample slice alignments. Patient 1 and 3 and patient 2 and 4 have 3 and 4 sections, respectively. (C) Mean accuracy for TOAST and DeST-OT across different cell types. (D) Execution time distribution for DeST-OT and TOAST. The violin plots illustrate the variability in computation time, while the black diamond markers indicate the median execution time, in seconds, for each method.

The results indicate that TOAST consistently outperforms DeST-OT across all patients, demonstrating its superior ability to establish biologically meaningful correspondences between aligned slices. TOAST also achieves higher accuracy for all but one cell type compared to DeST-OT (Figure 5 (C)). Finally, given the computational demands of large-scale spatial omics analyses, Figure 5 (D) shows the execution time distributions for both methods. These results, along with the findings in Figure S8, where we performed a controlled runtime analysis using simulated datasets of increasing size, indicate that TOAST not only improves alignment accuracy but also remains tractable even at tens of thousands of spatial locations, making it a scalable and efficient approach for spatial omics data integration.

### Spatial reconstruction

#### Spatial Mouse Atlas

Even though our OT methodology is not designed for the task of mapping spatial locations to single-cell data, we can apply it under the assumption of regional association of the gradient of expression to the spatial location which can be best observed in the context of early development ^30^. To this end, we performed semi-simulation experiments on the synthetic data generated from the Spatial Mouse Atlas datasets ^31^. Three different embryo slides were profiled with seqFISH ^4^ and single-cell (SC) gene expression profiles were provided along with the spatial coordinates of the cells. Following the approach in Hao et al. ^32^, we treated the gene expression data as pseudo-SC data without the spatial coordinates, and treated the spatial coordinates as the ground truth to test the predictions of our transport plan. We generated pseudo-spatial transcriptomics (ST) data to mimic key characteristics of the widely used 10X Visium platform ^2^: lower-than-single-cell resolution and partial coverage of cells within the tissue (Figure 6 (A)). The gene expression for each spot was calculated as the sum of the gene expression values of all the covered cells, while the true cell type proportions for each spot were determined based on the cell type annotations of the cells it encompassed. We compared TOAST to traditional FGW, DeST-OT, NovoSpaRc and Tangram. Given a transport plan obtained by any of the aforementioned methods, the spatial coordinates of a single cell were computed as the average of the coordinates of the Visium spots mapped to that cell, weighted by the transport probabilities. Mean Absolute Error (MAE) and Pearson Correlation Coefficient were used to measure the distance between the ground truth and predicted locations. Figure 6 (B) shows the results of this comparison. On both metrics, TOAST consistently outperforms the other methods, with DeST-OT and FGW reaching slightly lower performances. Tangram and NovoSpaRc show significantly worse spatial reconstruction abilities compared to the other methods. For each of three embryo slides, in Figure 6 (C), we show that the transport matrix produced by TOAST learns a mapping that strongly preserves the topology of the original single-cell space. Figure S5 shows this qualitative comparison for all the evaluated methods. As quantitatively indicated in Figure 6, FGW, DeST-OT and TOAST are better able to reconstruct the spatial relationships compared to Tangram and NovoSpaRc, which fail to preserve the spatial distribution of the single cells. Finally, we assessed the ability of TOAST to reconstruct the cell type relationships by computing a proximity-based enrichment score (see Methods) for each cell type in the original and reconstructed seqFISH data. For one of the samples, Figure 7 (A) shows that the reconstructed spatial cell type relationships closely resemble the ones in the original data. A high Spearman coefficient, Figure 7 (B), confirms this similarity. Figure S6 and Figure S7 depict the same analysis and similar results for the remaining samples.

**Figure 6:**
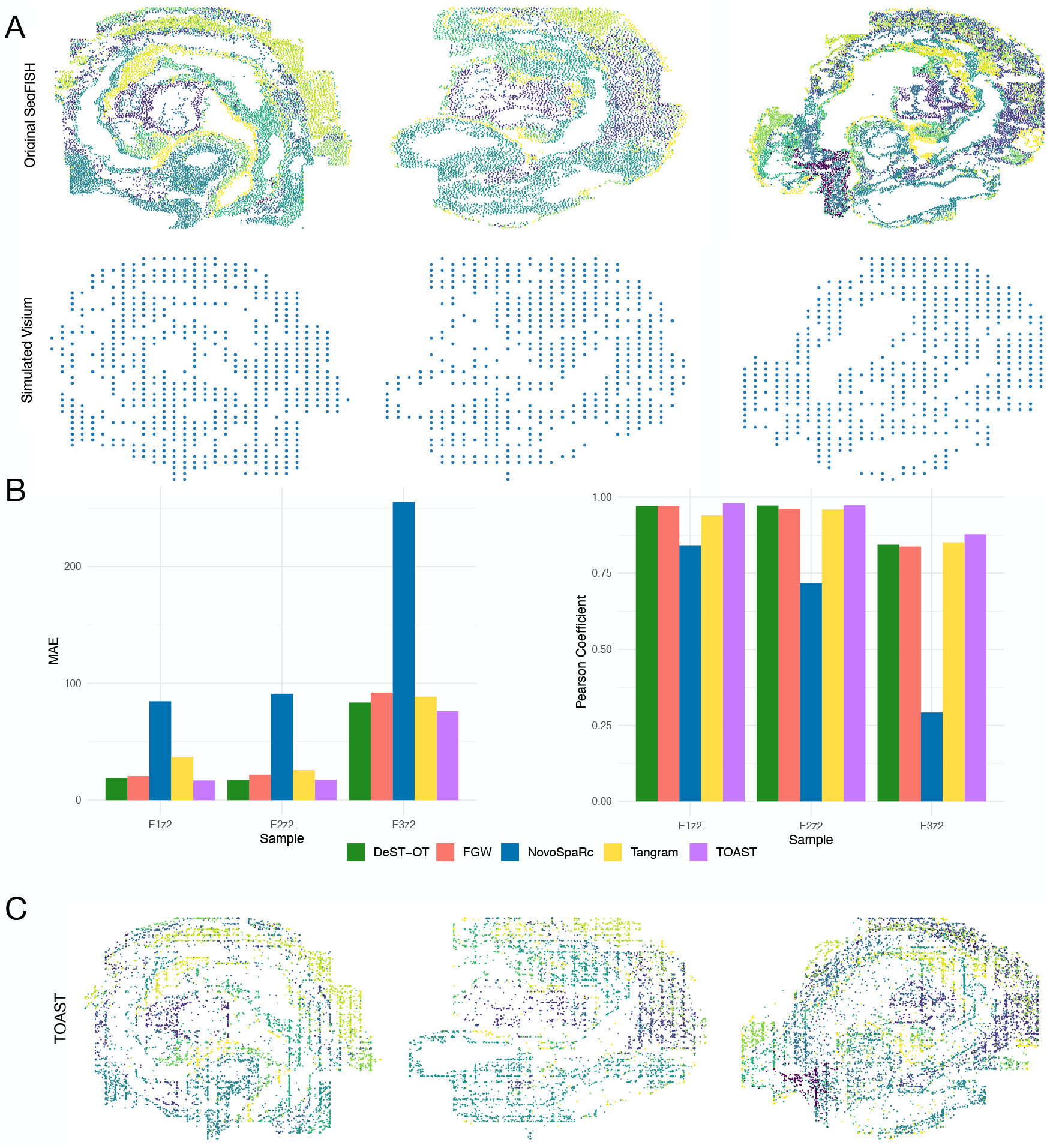
TOAST reconstructs spatial relationships in semi-simulated spatial transcriptomics data. (A) Original seqFISH spatial data and Simulated 10X Visium with lower resolution and coverage. (B) Mean Absolute Error (MAE) and Pearson correlation between ground truth and predicted locations for three embryo slides for FWG, TOAST, DeST-OT ^17^, Tangram ^27^ and NovoSparc ^28^. (C) The reconstructed spatial distribution by TOAST versus the ground truth spatial distribution. Colors indicate different cell types or regions.

**Figure 7:**
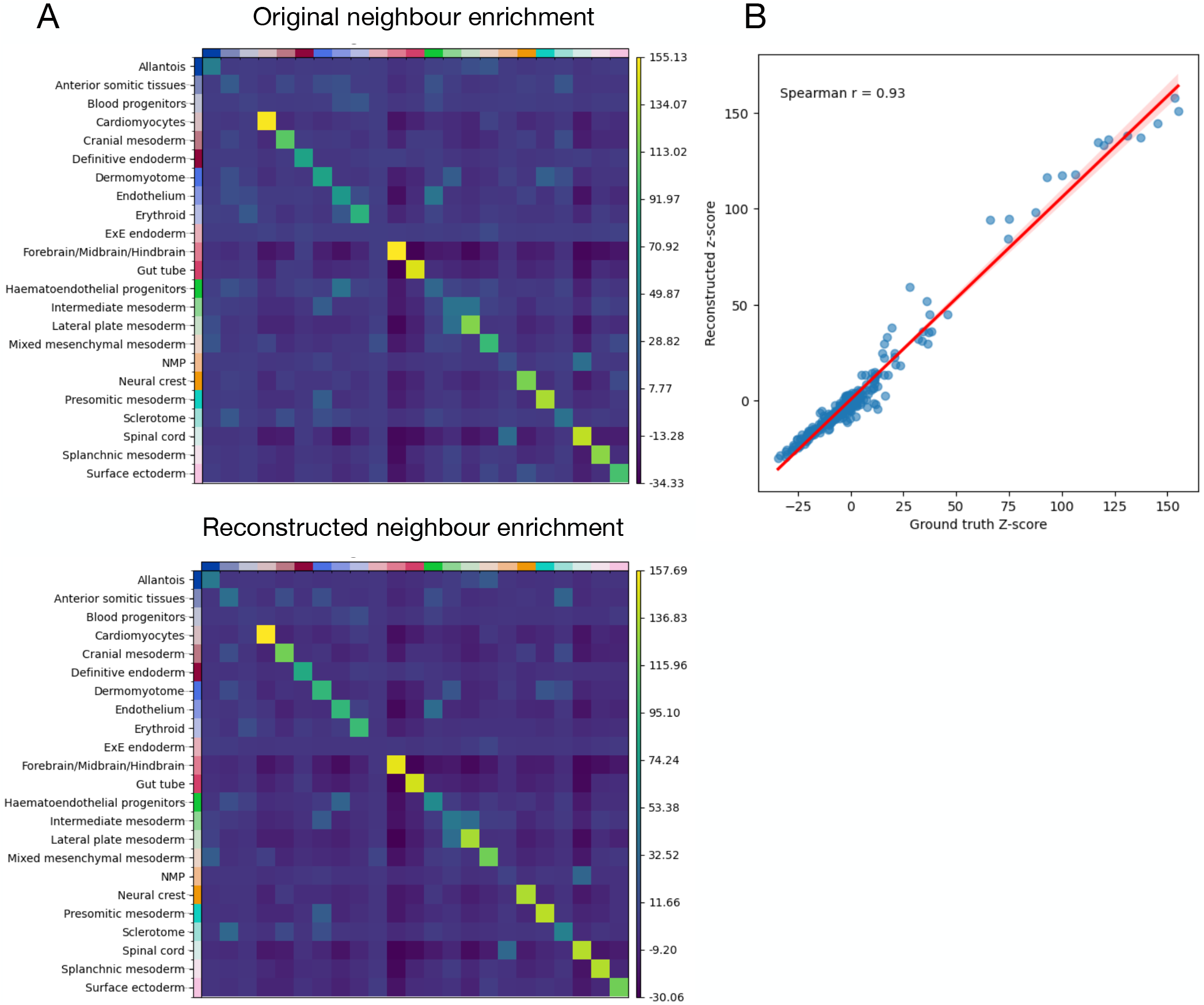
Spatial reconstruction performance of our topography aware OT framework on one sample from the Spatial Mouse Atlas. (A) Comparison of original and reconstructed neighbor enrichment for sample E1z1 from the Mouse Atlas ^31^. Heatmaps show the neighbor enrichment scores for different cell types in the original slice (top) and the reconstructed slice (bottom). The TOAST reconstructed enrichment matrix closely mirrors the original, demonstrating the model’s ability to preserve local tissue architecture. (B) The strong correlation between recon-structed and ground truth z-scores confirms the similarity between the original and reconstructed spatial interactions.

We performed a similar qualitative evaluation of our framework to analyze the tumor microenvironment in a human squamous cell carcinoma (hSCC) dataset ^33^. Figure S9 shows that TOAST is able to accurately reconstruct the cell type composition and the spatial relationships of the tumor microenvironment (see Supplementary Material).

## Discussion

Traditional OT-based alignment methods applied to omics data focus primarily on minimizing discrepancies between molecular profiles while neglecting the critical role of the spatial organization of tissues. The spatial arrangement and functional state of cells are not independent, but they are shaped by interactions with neighboring cells, extracellular matrix components, and local signaling interactions. Intra- and intercellular relationships give rise to spatial patterns that are characteristic of both physiological and pathological conditions. In this study, we introduced a scalable, topography aware Fused Gromov-Wasserstein optimal transport framework, TOAST, that explicitly incorporates spatial constraints into the classical FGW formulation. By integrating spatial coherence and neighborhood consistency in its objective function, our approach disentangles cell-intrinsic variability from the expression profiles in each cell’s neighborhood and effectively models molecular heterogeneity and tissue architecture. Extensive evaluation shows that TOAST leads to biologically meaningful mappings in spatial omics datasets.

On simulated data, our results demonstrate that the inclusion of spatial coherence and neighborhood consistency improves the alignment of both clustered and disorganized tissue regions, with traditional OT methods struggling to capture these nuanced spatial patterns, often leading to biologically implausible mappings. When applied to real spatial transcriptomics datasets, TOAST consistently outperforms competing methods across multiple evaluation criteria. In the human dorsolateral prefrontal cortex (DLPFC) dataset, our approach achieves the highest accuracy in aligning cortical slices, preserving the integrity of spatially distinct neuronal layers more effectively than traditional OT frameworks. Similarly, in the Axolotl brain dataset, TOAST successfully recovers developmental trajectories by accurately aligning cell states across consecutive time points and by modeling the local tissue microenvironment during the alignment process. Beyond spatial transcriptomics, we demonstrate the versatility of TOAST by applying it to Imaging Mass Cytometry (IMC) data from multiple cancer types. Our framework successfully aligns intra-sample sections, outperforming other methods in preserving the tissue spatial organization. Finally, our results on the mouse atlas dataset showcase the ability of TOAST to accurately reconstruct spatial organization across anatomical regions. By leveraging spatial constraints, TOAST effectively captures the underlying spatial structure of the mouse embryos, preserving tissue-specific arrangements and improving the biological relevance of cell-cell mappings. Beyond the aforementioned applications, our framework provides a principled way to align and integrate multi-omics spatial datasets and could have further implications for understanding developmental trajectories, disease progression, and tissue regeneration.

While our framework offers significant improvements over traditional OT-based methods, it is not without limitations. First, TOAST relies on a *balanced* optimal transport (OT) formulation, where the total mass transported from one spatial slice to another is constrained to be equal. While suitable for controlled datasets with stable cell type proportions, this assumption may not hold in biological systems where cell proliferation, death, or migration alter mass distributions. In such scenarios, an *unbalanced* OT formulation, which relaxes mass conservation constraints by incorporating additional regularization terms, could provide more flexible and meaningful mappings. Second, our method models pairwise alignment between spatial domains, which may limit its ability to capture more complex temporal trajectories or multi-sample correspondences. A possible extension would be to adopt a multimarginal OT framework, allowing simultaneous alignment of multiple spatial slices or time points. This could enable a more comprehensive modeling of dynamic processes such as development, regeneration, or disease progression. Looking forward, TOAST opens avenues for a range of downstream tasks beyond alignment of spatially resolved omics. For instance, it could be used to reconstruct spatial trajectories, annotate disorganized tissue regions, or infer changes in local tissue microenvironments across conditions or perturbations. In addition, future work could explore adaptive spatial regularization, where neighborhood constraints are dynamically tuned based on local spatial resolution or cell density, enhancing robustness across structurally diverse tissue architectures.

## Methods

### Topography Aware Fused Gromov-Wasserstein

The Fused Gromov-Wasserstein (FGW) distance is an optimal transport framework that aligns distributions based on both *feature similarity* and *structural similarity*. The usual formulation of FGW is

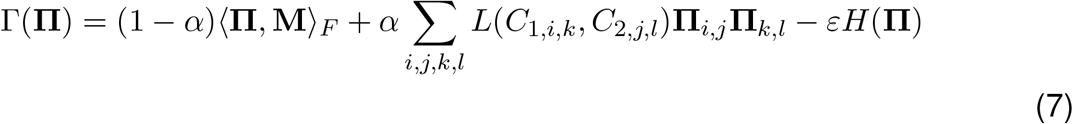

where *H*(**Π**) = ∑ _*i,j*_ **Π**_*i,j*_ log(**Π**_*i,j*_) is the entropy of the transport matrix and *ε >* 0 controls the strength of this entropic regularization. In this work, we extend FGW to

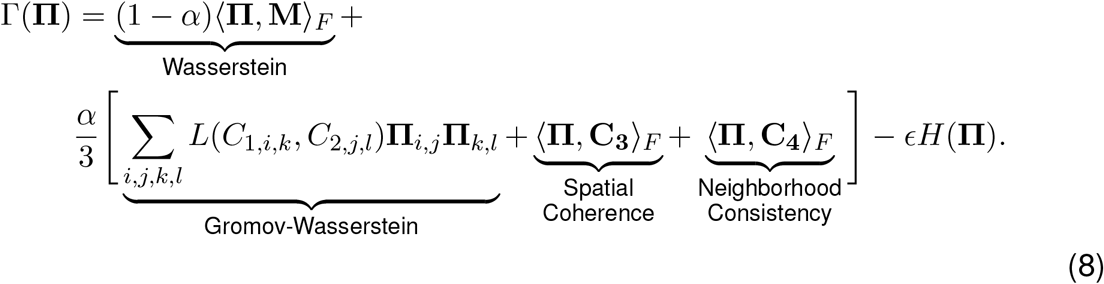

Given some cost matrix **M**, where **M**_*ij*_ represents the transportation cost between positions indexed by *i* and *j*, optimal transport seeks to find a coupling between two probability distributions. Let *p ∈ Δ*^*n*^ and *q ∈ Δ*^*m*^ be discrete probability distributions over the source and target spaces, respectively, where *p*_*i*_ and *q*_*j*_ denote the probability mass associated with locations indexed by *i* and *j*. Here, “^*n*^ represents the *n*-dimensional probability simplex defined as

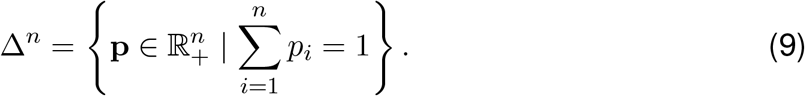

This ensures that the entries of *p* and *q* are non-negative and sum to one, making them valid probability distributions. Then, the solution of the entropically normalized optimal transport between *p* and *q* is the convex optimization problem

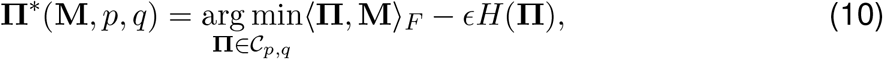

where 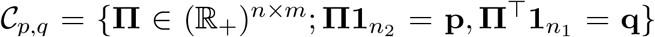 is the set of possible couplings between *p* and *q*. As shown in Cuturi ^26^, the solution is **Π**^*^ (**M**, *p, q*) = diag(*a*)*K*diag(*b*) where 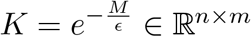 is the Gibbs kernel associated to **M** and (*a, b*) are computed as

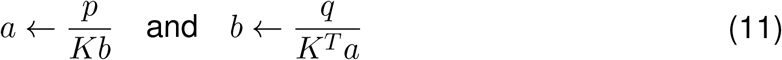

using Sinkhorn iterations ^34^. The Gromov-Wasserstein (GW) term measures the discrepancy between the two distance matrices **C**_**1**_ ∈ℝ^*n× n*^ and **C**_**2**_ ∈ ℝ^*m× m*^ defined on space *p* and *q*, respectively. For simplicity define

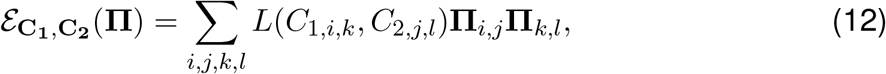

where *L* is some loss function to account for the discrepancy between the distance matrices. Using the 4-way tensor

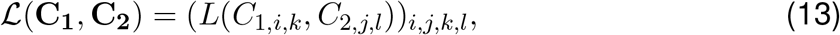

we have 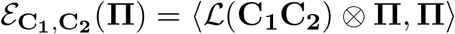 where ⊗ defines tensor-matrix multiplication. In Peyréet al. ^35^, the authors showed how to efficiently compute ℒ (**C**_**1**_**C**_**2**_) ⊗ **Π** for a general class of loss functions. We define the entropic approximation of the Gromov-Wasserstein formulation in Equation 8 as

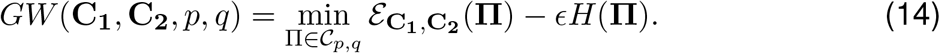

Equation 14 is a non-convex optimization problem that can be solved using projected gradient descent (PGD), where the projections are computed according to the Kullback-Leibler (KL) metric. An iteration of this algorithm reads as

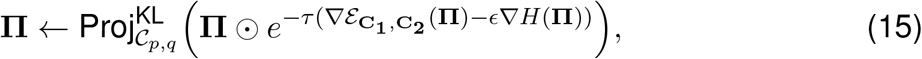

where τ is the step size, and the KL projection of any matrix *K* is given by:

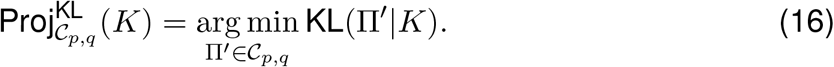

Essentially, Equation 16 projects the transport plan onto the feasible set using KL divergence, ensuring valid marginals while preserving entropy regularization. As shown in Peyréet al. ^35^, Benamou et al. ^36^, for small enough steps of τ, the projection is just the solution to the regularized transport problem in Equation 10. Hence, minimizing the problem in Equation 8, reduces to the following Sinkhorn iterations:

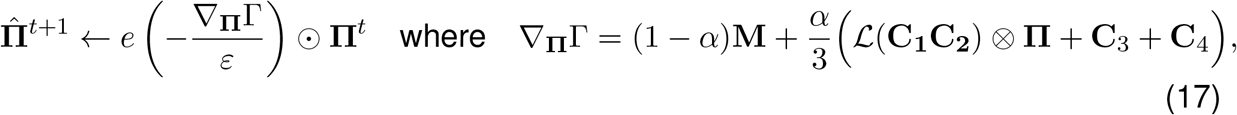

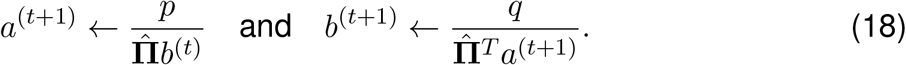

The transport plan is iteratively computed from Equations 17 and 18 as

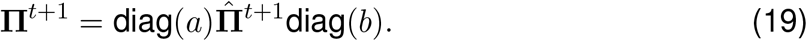

### Distance functions

For our topography aware Fused-Gromov Wasserstein formulation in Equation 5, we define multiple loss functions, operating in the principal component space ℤ^*p*^, to compute distances between cells in the source and target domains. In detail, the cost matrix **M** is computed as

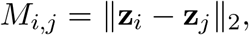

where **z**_*i*_ and **z**_*j*_ are the PCA embeddings for a point in the source and target domain, respectively. The matrix **C**_**3**_, enforcing spatial coherence, is computed as

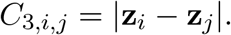

Similarly, **C**_**4**_, designed to model neighborhood consistency, is given by:

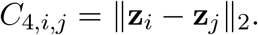

The matrices *C*_1,*i,k*_ and *C*_2,*j,l*_, which define structural relationships between points in the source and target domains, are computed as

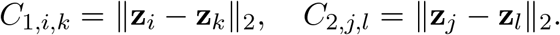

Finally, the function *L*, capturing pairwise dependencies between transported distributions, is defined as

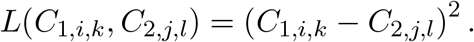

### One-dimensional simulation

To simulate the one-dimensional spatial transcriptomics data, we followed the approach described in Halmos et al. ^17^. We considered a pair of one-dimensional tissue slices, each consisting of 101 spatially ordered spots, spanning from −*N* to *N*, where *N* = 50. The spatial coordinates for both slices are represented by:

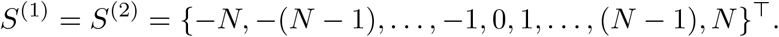

The spots are organized in clustered or disorganized regions: clustered regions only contain cell type A or cell type B, while disorganized regions exhibit alternation of the two cell types. The partition for slice 1 is given by:

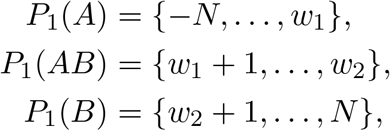

where *w*_1_ = −20 and *w*_2_ = 20 mark the boundaries of the clustered and disorganized regions. Similarly, the partition for slice 2 is:

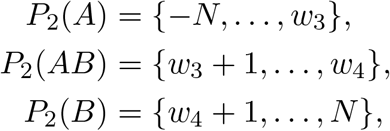

where *w*_3_ = −15 and *w*_4_ = 10 define the boundaries in the second slice.

Random vectors **v**_1_, **v**_2_, **v**_3_, **v**_4_, independently and uniformly sampled from the unit sphere in ℝ^4^, are used to create orthogonal feature gradients for cell types A and B. The features are constructed as

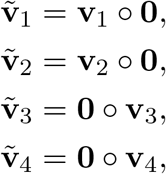

where **0** ∈ ℝ^4^ is the zero vector, and denotes vector concatenation.

Features are then generated by linearly interpolating between two fixed vectors:

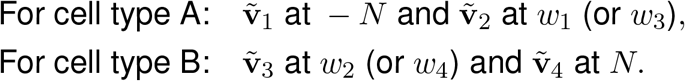

After generating the features, Gaussian noise is added to simulate variability. The noise is sampled from a normal distribution with mean *µ* = 0 and standard deviation σ= 0.1. This setup ensures that features for cell types A and B remained orthogonal, with each region exhibiting distinct gradients along four-dimensional feature spaces, with added noise reflecting natural variability.

### Two-dimensional simulation

To extend the one-dimensional model to two-dimensions, we define a centered, circular ellipse *E* of radius *r* = 20, represented as

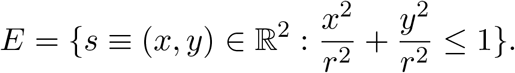

Next, we define *T*, a subset of the integer square lattice ℤ^2^, consisting of all integer pairs whose sum is odd:

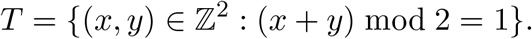

This results in a triangular lattice structure, which mimics the spatial organization of spatial transcriptomics data produced by platforms like 10X Genomics Visium ^2^. The spatial coordinates for both tissue slices, denoted *S*^(1)^ = *S*^(2)^ = *S*, are given by the intersection *E* ⋂ *T*, representing spots within the circular ellipse. To partition each slice into clustered and disorganized regions, we define pivot lines for the *y*-coordinates. For slice 1, the pivot lines are 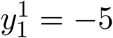 and 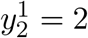. For slice 2, the pivot lines are 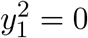 and 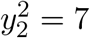. Based on these pivots, we define three areas for slice 1:

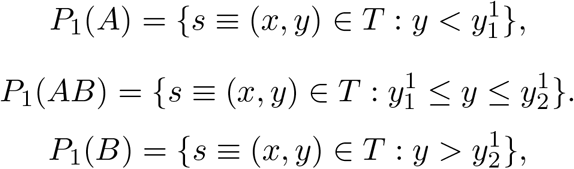

For slice 2, the partitions follow a similar structure:

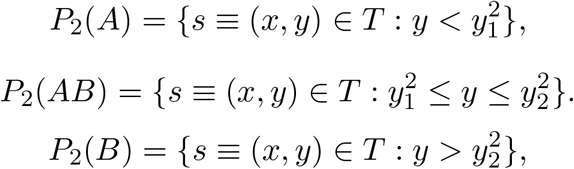

For each slice *i*, let *R*_*i*_(*A*), *R*_*i*_(*B*) and *R*_*i*_(*AB*) denote the smallest bounding rectangles for the clustered and disorganized regions *P*_*i*_(*A*), *P*_*i*_(*B*) and *P*_*i*_(*AB*) respectively. For feature assignment, we use the first four unit vectors in the standard basis of ℝ^8^, denoted **e**_1_, **e**_2_, **e**_3_, **e**_4_, assigned to cell type A, and the last four vectors, **e**_5_, **e**_6_, **e**_7_, **e**_8_, assigned to cell type B.

At each spot *s* = (*x, y*) within a bounding rectangle *R* = [*x*_min_, *x*_max_] × [*y*_min_, *y*_max_], the feature vector is determined as a convex combination of the features decorating the top and bottom sides of *R*, plus the features decorating the left and right sides of *R*. The coefficients *λ*_*x*_ and *λ*_*y*_ are defined as

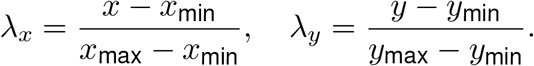

For cell type A, the feature vector is:

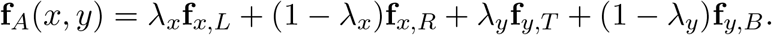

For cell type B, the feature vector is:

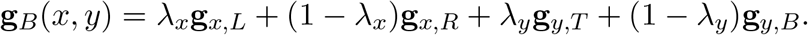

To simulate variability, Gaussian noise, sampled from a normal distribution with mean *µ* = 0 and standard deviation σ = 0.1, was added to the features.

### Evaluation metrics

For all comparisons to the related approaches, we computed an accuracy score based on the maximum probability assignment for each cell. In detail, for each cell *i* in the source domain, we identified its most probable match in the target domain given a pairwise alignment matrix **Π** = [π _*i,j*_] generated by a method as

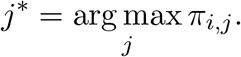

Using this assignment, we defined the accuracy as the proportion of correctly matched cells, where correctness was evaluated based on known biological annotations (for example, cell types). To explore the ability of the models to reconstruct the local spatial heterogeneity, we also quantified the discrepancy of cell type distributions between source and aligned slices using the Jensen-Shannon Divergence (JSD). Given two probability distributions *P* and *Q*, JSD is defined as

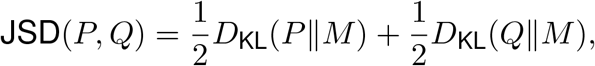

where 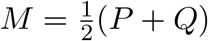 is the average distribution, and *D*_KL_(*P* ‖ *Q*) denotes the Kullback-Leibler (KL) divergence:

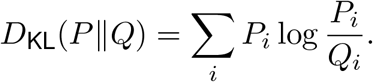

For each pair of aligned cells, we quantified the JSD of their nearest 20 neighbors, capturing local discrepancies in cell type distributions. The overall JSD score of a method was then computed as the mean JSD across all aligned cells:

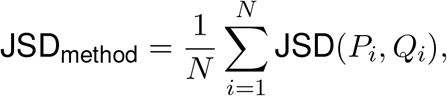

where *N* is the total number of aligned cells, and JSD(*P*_*i*_, *Q*_*i*_) represents the JSD between the local cell type distributions of the *i*-th aligned cell in the source and target domains. A lower overall JSD score indicates that the method preserves the local spatial organization of cell types more effectively. For the spatial reconstruction experiments on the Spatial Mouse Atlas, given the continuous nature of the predictions (spatial locations), we compared the models using Mean Absolute Error (MAE):

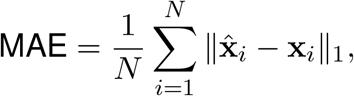

where 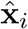 and **x**_*i*_ represent the predicted and ground truth spatial coordinates of cell *i*, respectively, and ‖ · ‖_1_ denotes the *L*_1_-norm. Additionally, we computed the Pearson correlation coefficient to assess the linear relationship between predicted and true spatial locations:

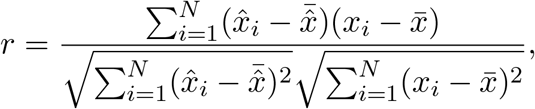

where 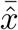 and 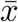 are the means of the predicted and ground truth coordinates, respectively.

### Related approaches

#### DeST-OT

Developmental SpatioTemporal Optimal Transport (DeST-OT) ^17^ is a semi-relaxed optimal transport method introduced to model cellular growth, death, and differentiation processes. DeST-OT generates mapping probabilities for each cell in one slice to each cell in the other slice. Therefore, we selected the positions with the maximum probability as the prediction in our comparison experiments. We evaluated DeST-OT using the default parameters suggested by the authors, but setting the optimization problem as a balanced one to fairly compare to the other methods.

#### NovoSpaRc

NovoSpaRc ^28^ is an optimal transport method which maps cells to tissue locations. The predefined locations for the mapping were constructed based on the reference ST dataset. Following the NovoSpaRc pipeline, we first identified marker genes from the reference dataset and then performed the mapping process based on these markers. We run NovoSpaRc using the default parameters suggested by the authors.

#### Tangram

Tangram ^27^ is a deep learning framework that aligns single-cell and single-nucleus RNA-seq data to various forms of spatial data collected from the same region. Similarly to the other methods, Tangram generates mapping probabilities for each cell. We run Tangram using the default parameters suggested by the authors for 200 epochs.

#### Paste2

Paste ^15^ was originally introduced as a balanced optimal transport method to align and integrate ST data from multiple adjacent tissue slices. Paste2^16^ was introduced as an extension to Paste as a method for partial alignment of slices. Paste2 is based on the Fused Gromov-Wasserstein Optimal Transport problem. We run Paste2 with the default parameters suggested by the authors but setting the slice overlap argument to 1 (all mass is transported from the source slice to the target slice).

#### FGW

The traditional Fused Gromov-Wasserstein formulation was implemented in Python using the POT package ^37^.

### Spatial coherence and neighborhood consistency

To compute the spatial coherence and neighborhood consistency matrices in Equation 5, we build a *k*-nearest neighbor graph from the (*x, y*) coordinates of each slice. Therefore, for each cell, the spatial coherence and neighborhood consistency values are simply defined as in Equation 2 and Equation 3, respectively. We set the value of *k* to 10 for the Spatial Mouse Atlas and Human Dorsolateral Prefron Cortex dataset, and 5 for the Axolotl Brain Stereo-seq dataset. On the latter, given the lack of spatial coordinated for the single-cell (SC) data, we computed the spatial coherence scores and neighborhood averages by constructing a *k*-nearest neighbors graph based on the SC expression distances rather than spatial coordinates.

### Data preprocessing

We preprocessed the data using the Scanpy Python package ^38^. Specifically, we performed library size normalization (sc.pp.normalize total) followed by log-transformation (sc.pp.log1p). Principal component analysis (PCA) was then applied to reduce dimensionality.

### Neighborhood enrichment

We performed neighborhood enrichment on the cell types from the Spatial Mouse Atlas using nhood enrichment from the Squidpy Python package ^39^. This function computes a neighborhood enrichment z-score for each cell type by a permutation test. We used the function’s default parameters.

## Data and code availability

This study used publicly available data and the original data can be obtained at the following links: (1) 10x Visium human dorsolateral prefrontal cortex from https://research.libd.org/spatialLIBD/, (2) Axolotl brain Stereo-seq from https://ftp.cngb.org/pub/SciRAID/stomics/STDS0000056/stomics/, (3) seqFISH data on mouse organo-genesis from https://crukci.shinyapps.io/SpatialMouseAtlas/, (4) the Imaging Mass Cytometry data from the Integrated iMMUnoprofiling of large adaptive CANcer patient cohorts ^29^ (IMMUcan) was accessed through the Bioconductor package `imcdatasets`. The source code is accessible at https://github.com/cecca46/TOAST. TOAST is implemented in Python and can be seamlessly integrated into pipelines using Scanpy ^38^ and Squidpy ^39^, facilitating downstream analysis of spatial omics data.

## Acknowledgments

We gratefully acknowledge the contribution of Simon Gutwien for his assistance in the preparation of the figures presented in this work. For the purpose of open access, the author has applied a Creative Commons Attribution (CC BY) license to any Author Accepted Manuscript version arising from this submission.

