## Supplementary Data for "Topography Aware Optimal Transport for Alignment of Spatial Omics Data"

### A Supplementary figures

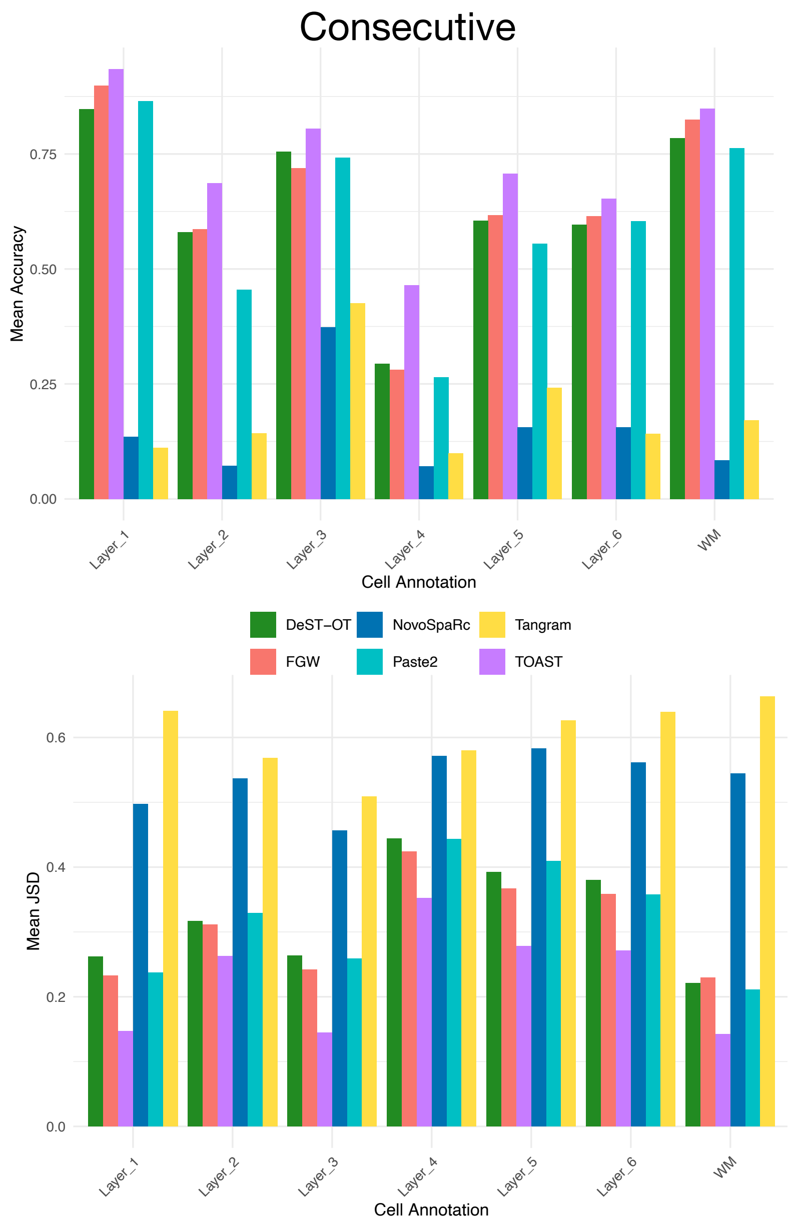

**Figure S1:** Quantitative comparison of pairwise alignment across all consecutive slices for FGW, TOAST, DeST-OT, Tangram, Paste2, and NovoSparc, evaluated in terms of accuracy and Jensen-Shannon divergence (JSD) for each annotated cell type.

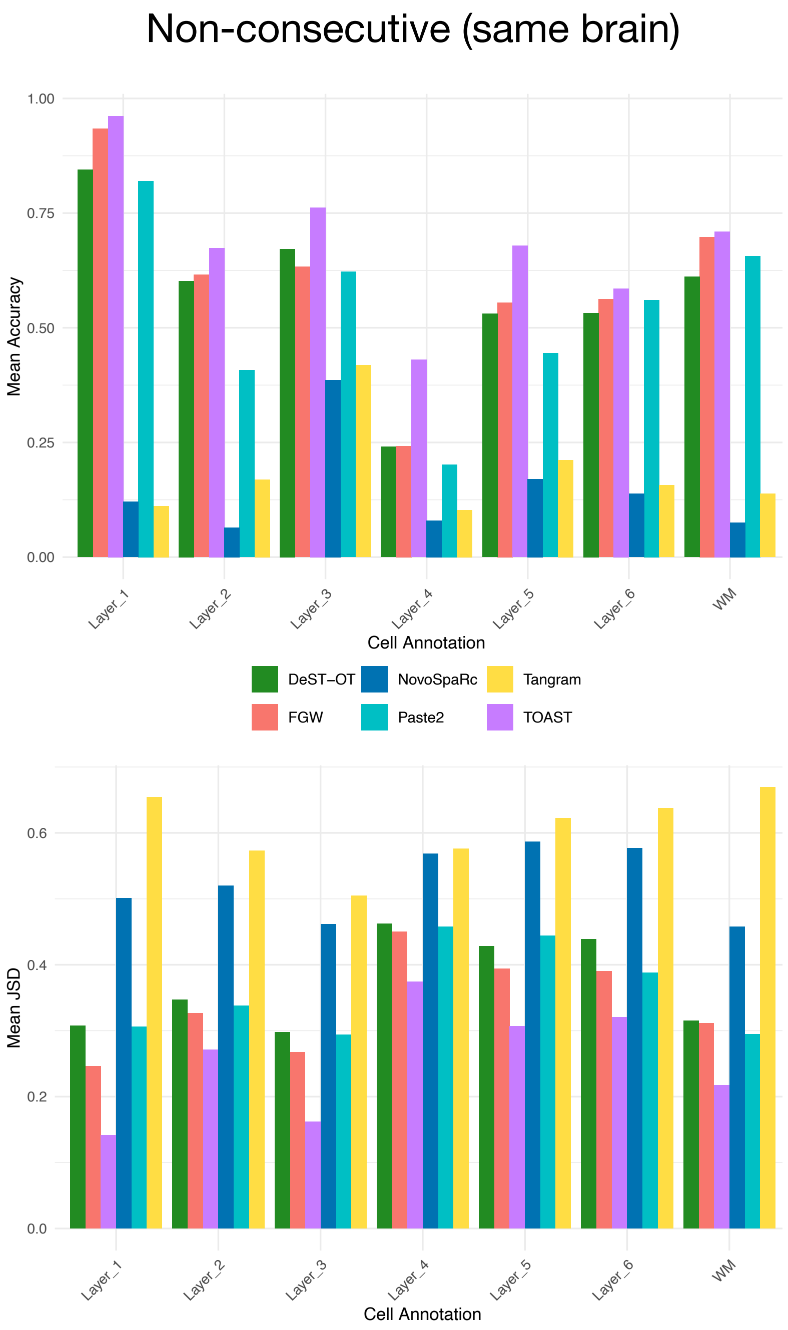

**Figure S2:** Quantitative comparison of pairwise alignment across all non-consecutive slices from the same brain for FGW, TOAST, DeST-OT, Tangram, Paste2, and NovoSparc, evaluated in terms of accuracy and Jensen-Shannon divergence (JSD) for each annotated cell type.

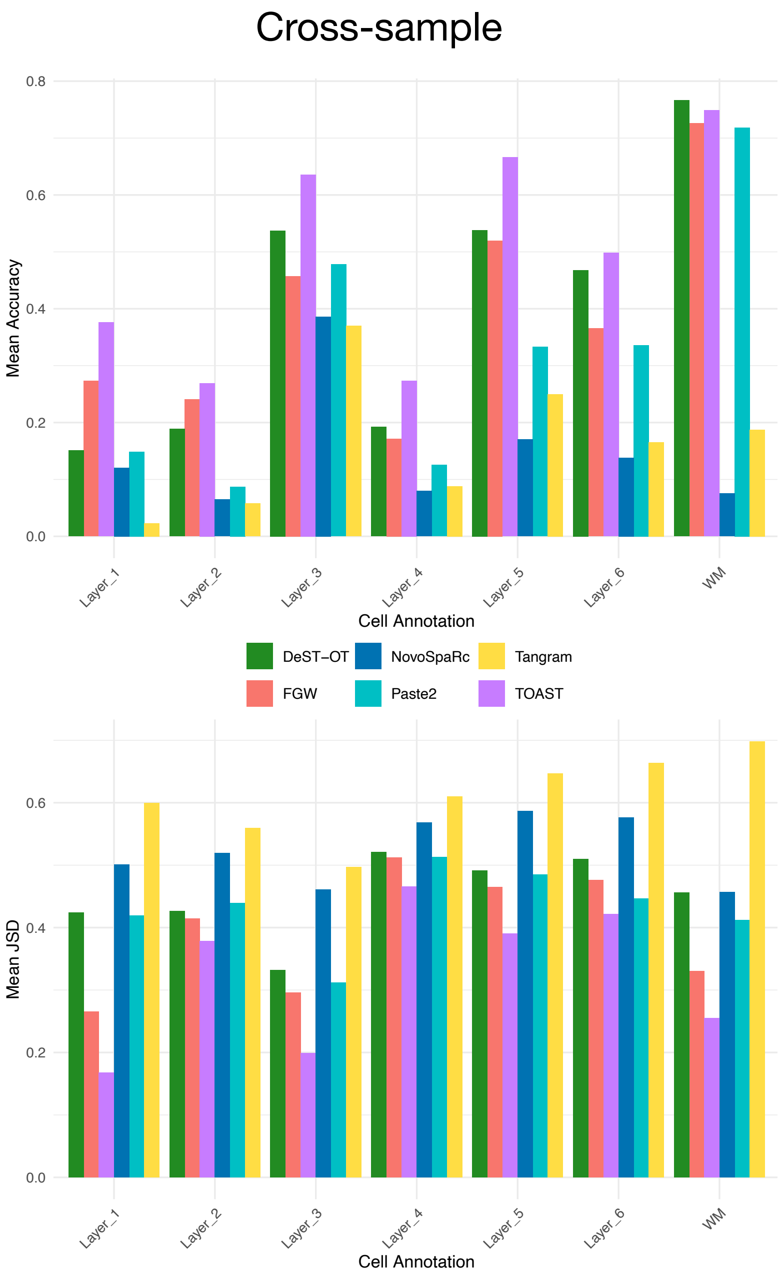

**Figure S3:** Quantitative comparison of pairwise cross-sample alignment for FGW, TOAST, DeST-OT, Tangram, Paste2, and NovoSparc, evaluated in terms of accuracy and Jensen-Shannon divergence (JSD) for each annotated cell type.

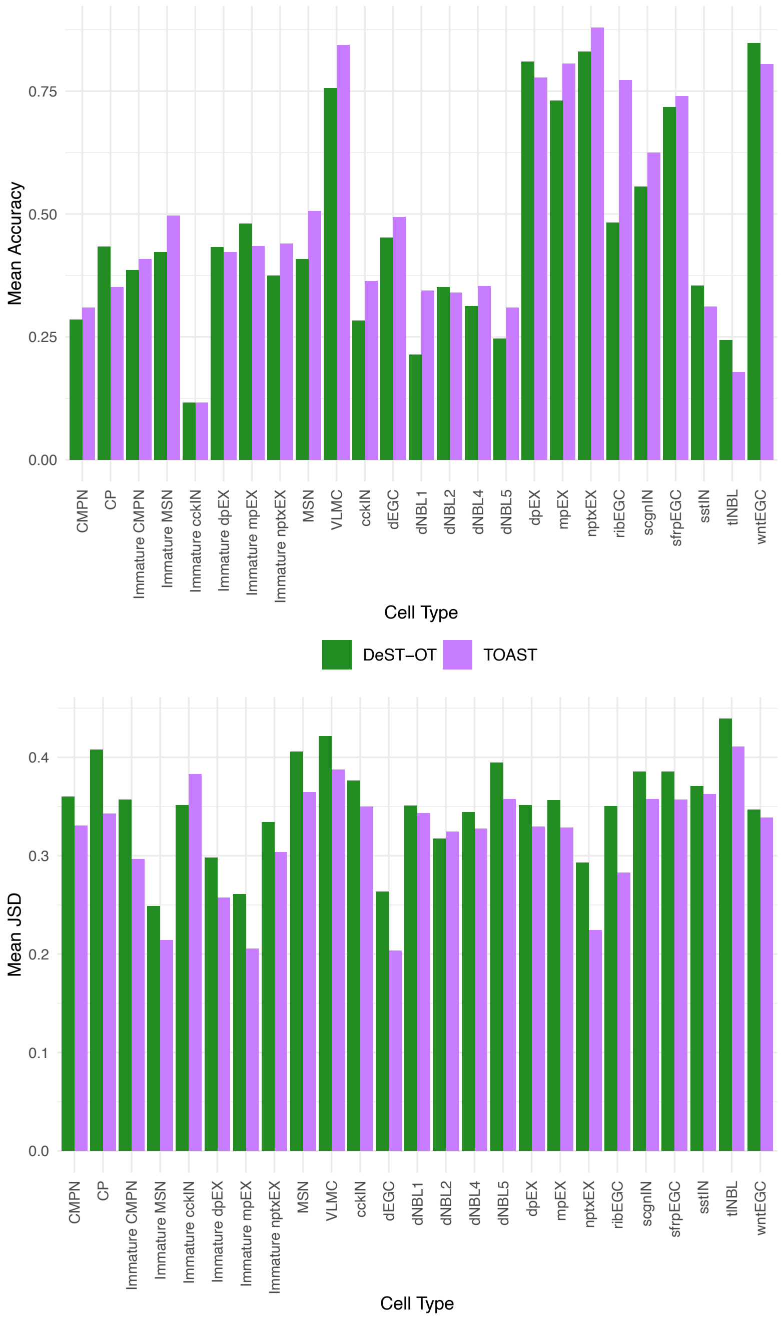

**Figure S4:** Quantitative comparison of pairwise alignment across all consecutive slices for TOAST and DeST-OT evaluated in terms of accuracy and Jensen-Shannon divergence (JSD) for each annotated cell type.

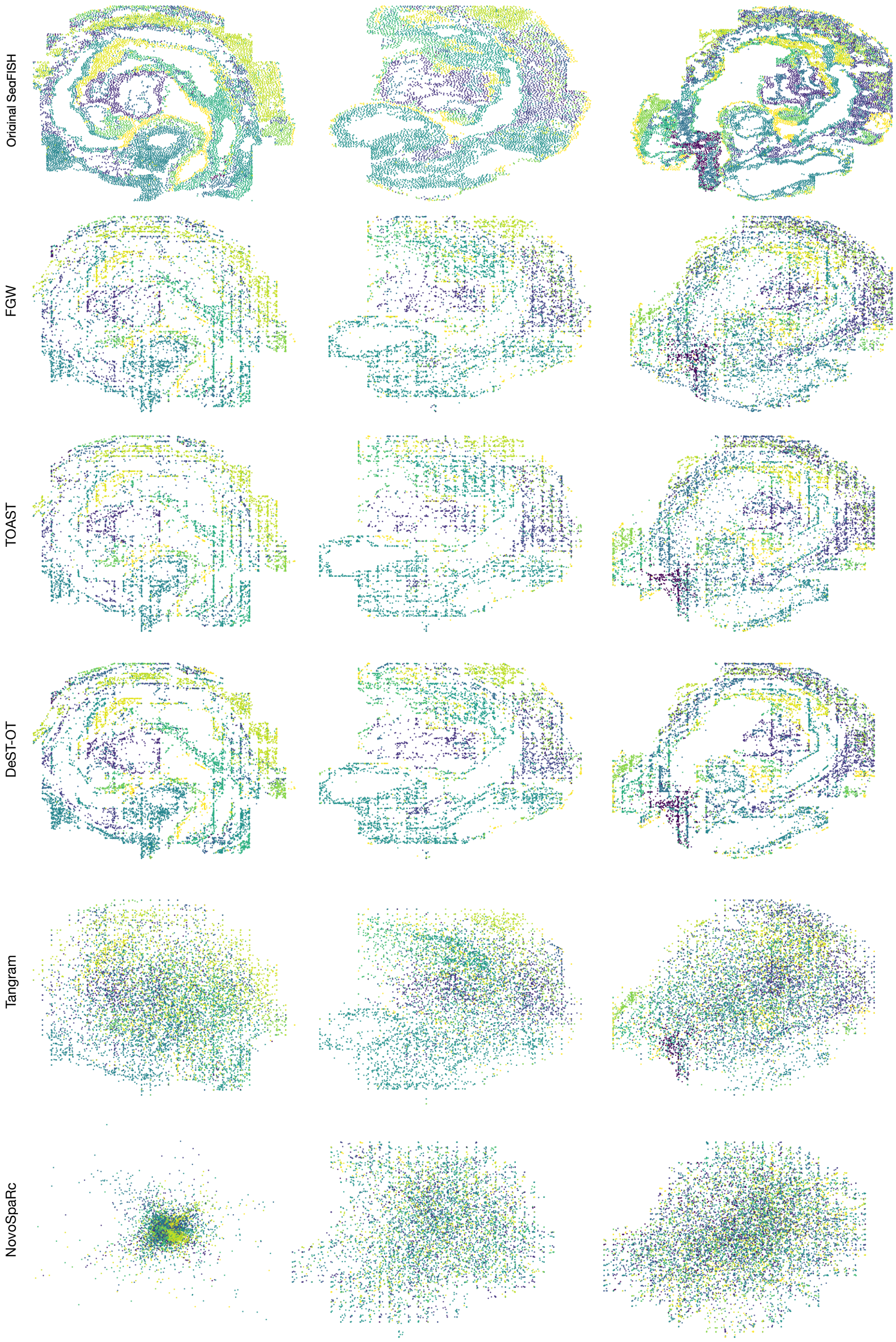

**Figure S5:** The reconstructed spatial distribution by FWG, TOAST, DeST-OT, Tangram and NovoSparc versus the ground truth spatial distribution. Colors indicate different cell types or regions.

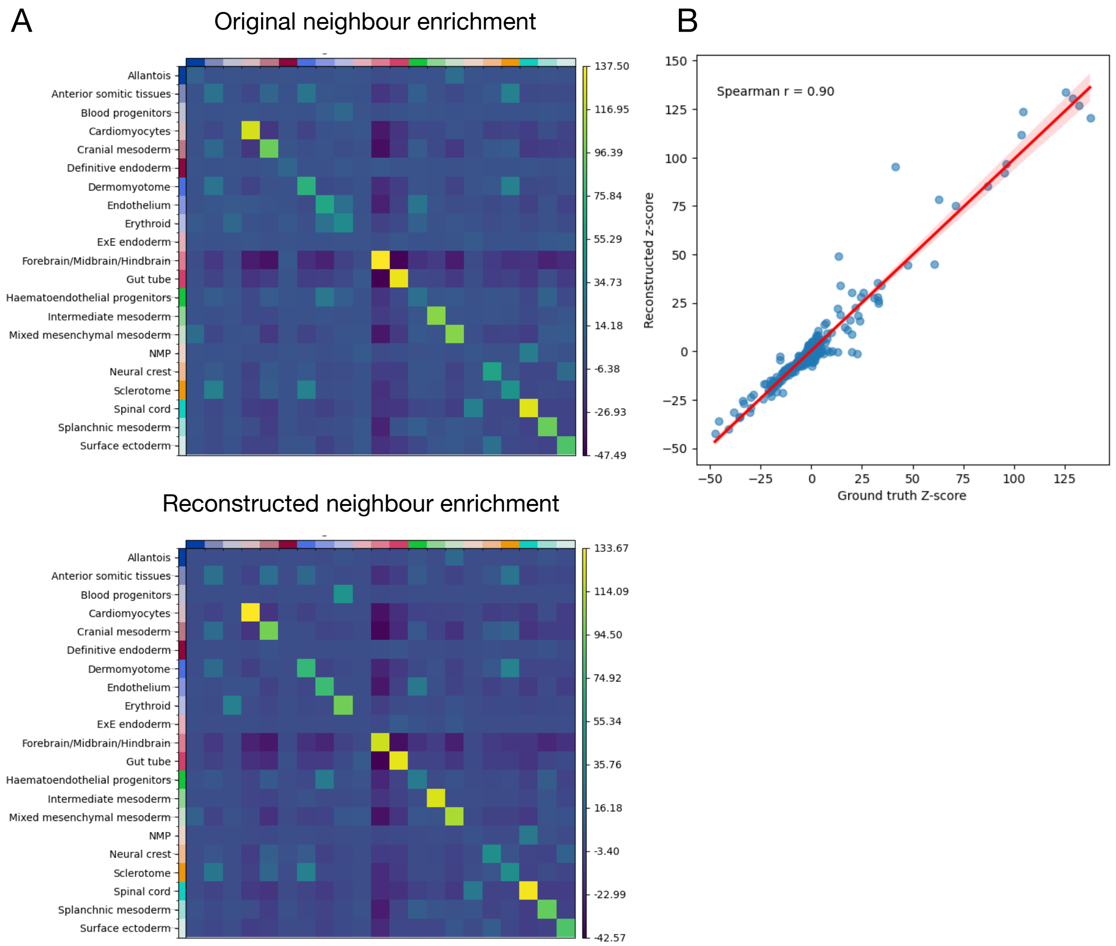

**Figure S6:** (A) Comparison of original and reconstructed neighbor enrichment for sample E1z2 from the Mouse Atlas. Heatmaps show the neighbor enrichment scores for different cell types in the original slice (top) and the reconstructed slice (bottom). (B) Correlation between reconstructed and ground truth z-scores.

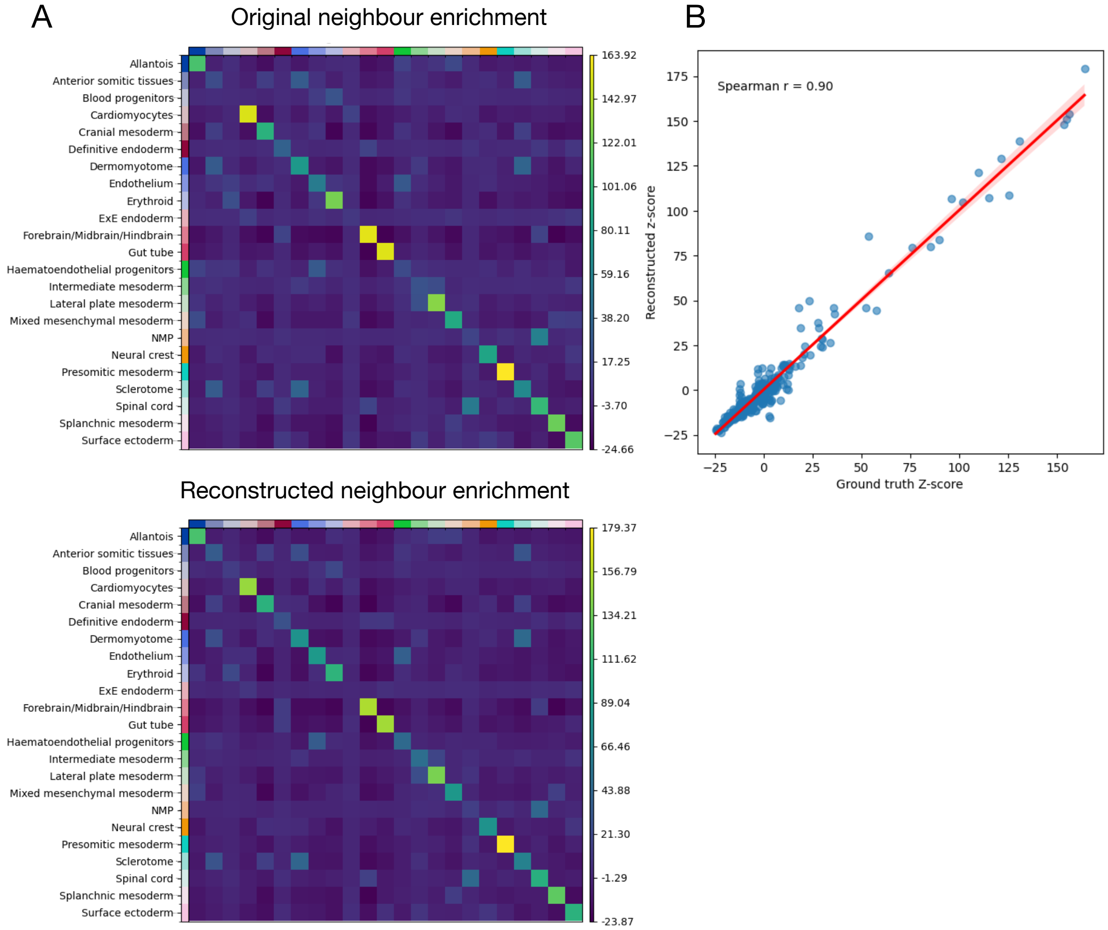

**Figure S7:** (A) Comparison of original and reconstructed neighbor enrichment for sample E1z3 from the Mouse Atlas. Heatmaps show the neighbor enrichment scores for different cell types in the original slice (top) and the reconstructed slice (bottom). (B) Correlation between reconstructed and ground truth z-scores.

#### B Additional experiments

**Scalability with Increasing Dataset Size** To evaluate the computational scalability of TOAST, we simulated two-dimensional spatial transcriptomics datasets with increasing numbers of spatial spots per slice, ranging from 1,500 to 19,000. These datasets retained spatial structure and molecular heterogeneity by preserving clustered and disorganized regions (see Methods). For each dataset, we measured the wall-clock time required to compute the transport plan using TOAST's topography aware objective. Figure S8 shows the percentage increase in runtime as a function of the number of spots per slice. Consistently with the nature of optimal transport computations, the trend in the plot suggests that the runtime grows approximately with the square of the number of spots, indicating a quadratic time complexity. Empirically, we observed that the runtime remains under one minute for datasets with fewer than 5,000 spots and increases to approximately 7.5 minutes for datasets with nearly 20,000 spots. This trend demonstrates that TOAST is well-suited and computationally tractable for analyzing high-resolution spatial datasets. All runtime benchmarks were performed on a Linux server running Ubuntu 20.04.6 LTS, equipped with two Intel(R) Xeon(R) Platinum 8260 CPUs (2.40 GHz, 48 cores, 96 threads total). All timing results were obtained using a single CPU core, without GPU acceleration.

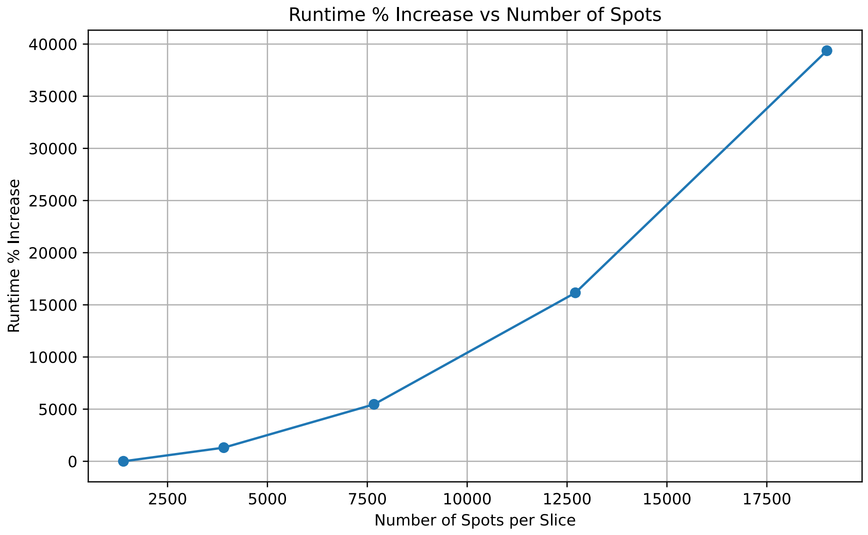

**Figure S8:** Percentage increase in runtime (seconds) as a function of the number of spots per slice.

**Human squamous cell carcinoma** The characterization of the tumor microenvironment is critical in all phases of cancer development and metastasis[40]. In these experiments, we applied our topography aware OT framework to a human squamous cell carcinoma (hSCC) dataset[33]. We used paired single-cell (SC) and spatial transcriptomics (ST) data from the hSCC tissue of a donor. The SC data was manually annotated, while the unannotated ST data were obtained by an early version of the 10X Visium. Here, we aimed at mapping the SC cell types onto the ST transcriptomics data. Given the lack of cell type annotation for the ST transcriptomics data, we computed the spatial coherence term using the cell cluster labels generated by running the Leiden algorithm[41]. The mixture of cell types of a Visium spot was computed as the average of the cell types of the single-cells mapped to that spot, weighted by the transport probabilities. Given the lack of ground truth annotations, we performed a purely qualitative comparison between the transport plans obtained from FGW and FGW with spatial coherence and neighborhood consistency (TOAST). We first verified that the tumor specific keratinocytes (TSKs) were colocalized with endothelial cells at the top tumor leading edge as described in the original study. From Figure S9 (A), it is clear that while both FGW and TOAST identified tumor specific keratinocytes, FGW fails to identify the enrichment of endothelial cells closer to the tumor region (green circle in Figure S9 (A)). We also explored the spatial organization of other keratinocyte (KC) subtypes identified by TOAST, including tumor basal, tumor cycling, and tumor differentiating KCs. Neighborhood enrichment analysis showed that TSKs tended to self-aggregate and spatially separate from other KCs (Figure S9 (B)). These findings, consistent with those of the original study, are crucial in revealing the spatial characteristics of the tumor microenvironment.

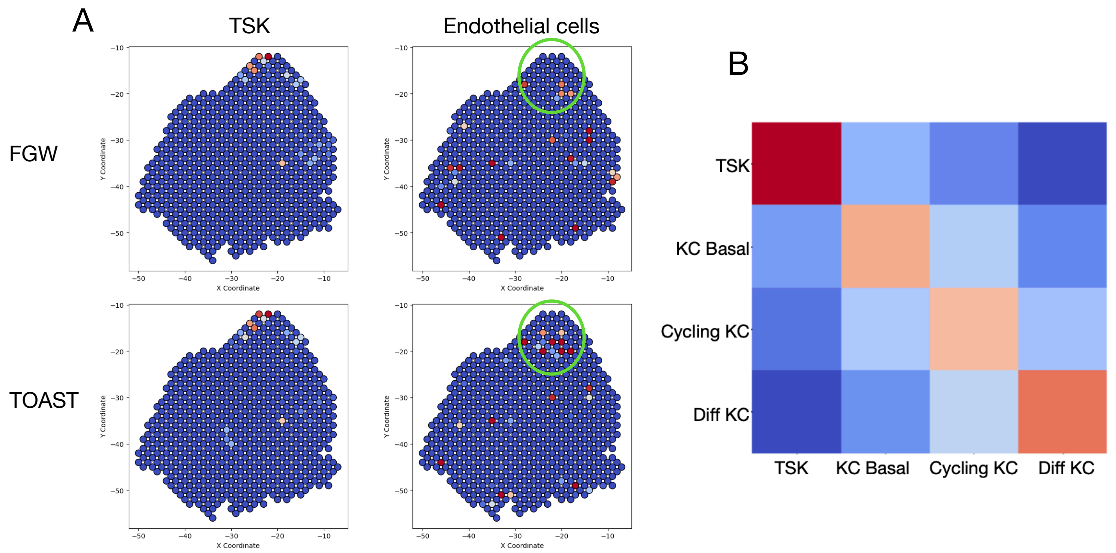

**Figure S9:** (A) Cell type distribution reconstruction by FGW and TOAST for one donor from the human squamous cell carcinoma (hSCC) dataset. While both FGW and TOAST identified tumor specific keratinocytes, FGW fails to identify the enrichment of endothelial cells closer to the tumor region described in the original publication. (B) Neighbor enrichment analysis on tumor keratinocyte subtypes. Red indicates a higher neighbor score.
